# A novel model demonstrating that human immune cells promote multiorgan SARS-CoV-2 dissemination and human T cells limit anti-viral innate immunity

**DOI:** 10.64898/2026.06.18.733232

**Authors:** Amber Wolabaugh, Vrushali V. Agashe, Zicheng Wang, Nichole M. Danzl, Christopher A. Parks, Mohsen Khosravi-Maharlooei, Xiaolan Ding, Hao Wei Li, Candace Castagna, Giorgia Zanetti, Katherine D. Long, Yasmeen S. Saad, Anjali Saqi, Moriya Tsuji, Megan Sykes

**Author notes:** Corresponding Author: Megan Sykes, Columbia Center for Translational Immunology, Columbia University College of Physicians and Surgeons, 650 West 168th Street, Suite 1512, New York, NY 10032. The authors have declared no conflict of interest.

## Abstract

Despite extensive studies, many questions remain regarding the pathology and mechanisms behind Coronavirus disease 2019 (COVID-19), and particularly on post-acute sequelae of COVID-19 (PASC). Because existing mouse models cannot recapitulate human immune responses, we developed a human immune system (HIS) mouse model with physiologic expression of hACE2. After intranasal infection, persistent viral RNA was observed in multiple organs for 8 weeks, despite the generation of human SARS-CoV-2-specific T cell responses. Human immune cells increased viral infection in the lung and non-pulmonary tissues. COVID-19-related pathology was recapitulated in the lungs, with fibrosis peaking at 14 days post-infection. Immune activation was detected in the lungs, hearts, intestines and brains of acutely infected mice and persisted at 8 weeks in the heart and lungs. The presence of human T cells correlated with attenuated innate antiviral transcriptional signatures in the lung, where a persisting cytotoxic mature CD4+ T cell population with JAK-STAT activation was enriched in infected mice. Thus, human T cells mount an antigen-specific but ultimately dysfunctional response, while paradoxically suppressing the innate interferon response, permitting chronic interferon activation and long-term viral persistence, recapitulating several immunologic and histopathologic features associated with PASC. This model will uniquely facilitate understanding of virus-human immune dynamics and therapeutic approaches to PASC.

## Introduction

Coronavirus disease 2019 (COVID-19) is a highly contagious respiratory disease caused by severe acute respiratory syndrome coronavirus 2 (SARS-CoV-2). Since its emergence at the end of 2019, it has caused over 700 million infections and claimed 7 million deaths worldwide, severely burdening global public health resources (1–4). Ongoing mutagenesis has given rise to continued outbreaks and the long-term effects of this disease are poorly understood. The risk of post-acute sequelae of COVID-19 (PASC) may increase with repeated infections (5, 6). COVID-19 manifestations are multisystemic and range from acute respiratory distress syndrome (ARDS) in acute infection, to fatigue, cognitive impairment, nausea, and dysautonomia in PASC (7). Despite a vast amount of research, many questions remain regarding the pathogenesis of COVID-19 and PASC. (8–10).

Studies of SARS-CoV-2 were initially limited by the lack of a high-throughput small animal model that could be used for evaluating human responses, as mice were not susceptible to infection by the original strain of virus due to genetic differences in the angiotensin-converting enzyme 2 (ACE2) receptor used to infect human cells (11–13). To overcome this, murine SARS-CoV-2 infection models often involved overexpression of hACE2 under common promoters such as the epithelial promoter cytokeratin 18 (K18). However, SARS-CoV-2 infection in this model often led to mortality from neurological pathologies, likely caused by the overexpression of hACE2 in the neuroepithelium compared to that in human COVID-19 patients (14–21). Even though subsequent variants of SARS-CoV-2 have become capable of infecting mice, murine innate and adaptive immune responses to infection harbor many differences from those of humans, such as those involving leukocyte subsets, T cell activation, antibody production, and costimulatory signaling (22–24). These differences continue to limit the ability of murine SARS-CoV-2 models to accurately reflect human infection. More recently, human immune system (HIS) mouse models have been developed to aid in the studies of human immune responses to SARS-CoV-2 infection. These models rely on the injection of hematopoietic stem cells (HSCs) into immunodeficient mice to reconstitute human immune cells, including T cells that develop in the murine thymic vestige (25, 26). However, human T cells that develop in a native mouse thymus do not undergo normal thymic selection and cause autoimmune disease, including enlargement of lymphoid organs and T cell infiltration into non-lymphoid tissues, which may interfere with the study of an infectious disease such as SARS-CoV-2 (27).

Furthermore, infection of HIS mouse models with COVID-19 remains dependent on the expression pattern of hACE2. While intranasal injection of SARS-CoV-2 into animals expressing respiratory hACE2 via Ad5- or AAV-hACE2 transduction systems (25, 26, 28) allows for effective modeling of immune pathology in the lungs, these models are incapable of representing the multi-organ disruption seen in human patients due to the lack of hACE2 expression in other targeted tissues (9).

To overcome the limitations of existing HIS mouse models of COVID-19, we produced a hACE2 knock-in, mACE2 knock-out NOD-scid-common gamma chain deficient (hACE2 KI NSG) mouse by replacing the murine ACE2 locus with hACE2 cDNA to allow for endogenous transcriptional regulation of hACE2 while disrupting expression of mACE2.

Upon removal of the native mouse thymus and implantation of human fetal thymus tissues and HSCs (29), we can then generate HIS mice capable of modeling SARS-CoV-2 pathogenesis in the context of a functioning human immune system with hACE2 expression at physiologic levels and in a tissue-specific pattern that closely resembles that of a human. This HIS mouse model demonstrates robust reconstitution of human T cells, B cells, and APCs (29–32). Additionally, the removal of the mouse thymus and inclusion of a human fetal thymus, in which both murine and human APCs are detected, effectively avoids autoimmunity and enables negative selection of human donor- and murine recipient-reactive T cells (27, 29, 33, 34). Mice reconstituting human B cells and myeloid cells, but lacking reconstituted T cells, can be generated by omitting only the human fetal thymus transplantation step in this process, thereby allowing us to pinpoint the role of human T cells in driving or controlling COVID-19 disease. Using this new hACE2 KI HIS mouse model, we performed acute and long-term SARS-CoV-2 infections to investigate viral dissemination, viral clearance, and virus-induced pathology across multiple tissues and dissected the impact of T cell immunity on these responses.

## Results

### hACE2 KI NSG mice demonstrate physiologic expression of hACE2

hACE2 CRISPR knock-in mice were generated on an NSG background to allow for subsequent human tissue engraftment and immune reconstitution. Insertion of the hACE2 cDNA sequence after the murine ACE2 start codon ensured physiologic expression of hACE2 while deleting the murine counterpart (Fig 1a). The hACE2 gene insert was validated at the genomic DNA level (Fig. 1b) and expression was characterized across fourteen different tissues through quantitative RT-PCR using hACE2-specific primers (Fig. 1c). hACE2 KI NSG mice had robust expression of hACE2, with murine lung tissue displaying transcript levels comparable to that of a human lung control. Of note, hACE2 expression was highest across all regions of the intestine, reproductive organs, liver, and spleen, with moderate expression in the kidney, heart and brain. As expected, hACE2 transcripts were undetectable in WT NSG mice (not shown). Immunohistochemical staining of hACE2 on formalin-fixed paraffin-embedded lung and intestinal tissue from hACE2 KI NSG mice revealed robust staining across bronchiolar epithelial cells and intestinal epithelial cells only in hACE2 KI and not WT NSG mice (Fig. 1d). Taken together, these data demonstrate both genetic and protein-level physiologic expression of hACE2, with an expression pattern resembling that described in humans (14, 35, 36).

**Figure 1:**
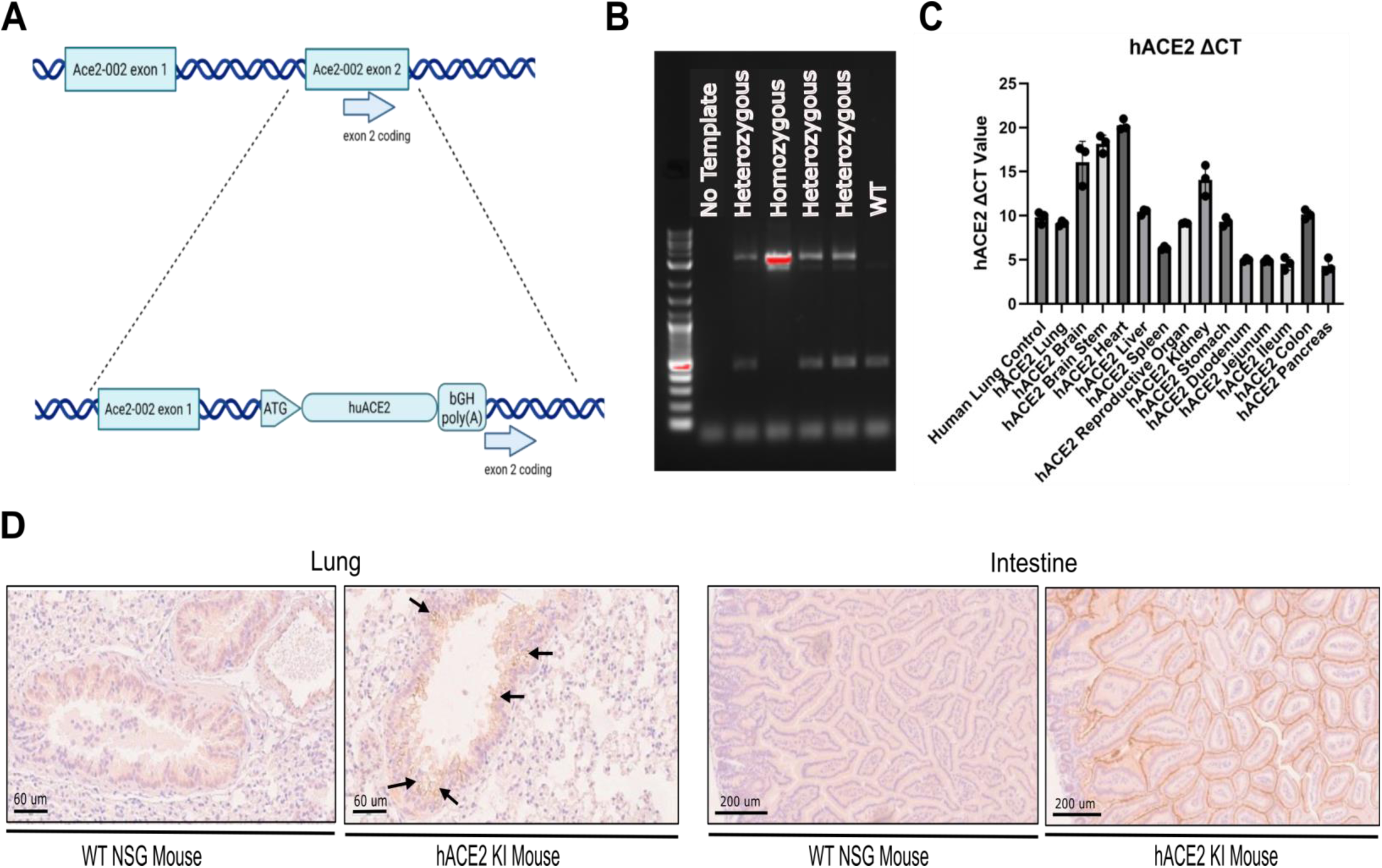
hACE2 KI NSG mice demonstrate physiologic expression of hACE2. **a.** Schematic of hACE2 gene knock-in under the murine ACE2 promoter, beginning in exon 2 of murine ACE2. **b.** Confirmation of the presence of the hACE2 gene (3081bp) in genomic DNA from hACE2 KI NSG mice, determined from hACE2-specific PCR and gel electrophoresis with a 1kb ladder. Murine ACE2 is 456bp and can be seen in heterozygous and WT mice, allowing for the differentiation of homozygous mice. **c.** Characterization of hACE2 expression across fourteen tissues in a homozygous hACE2 KI NSG mouse. ΔCT values were determined from a hACE2-specific one-step qPCR, with hACE2 expression normalized to murine GAPDH. Mouse samples were run alongside a human fetal lung control normalized to human GAPDH. n=3 **d.** Representative images of immunohistochemistry staining for hACE2 performed on formalin-fixed paraffin-embedded lung and intestinal tissue from both hACE2 KI NSG mice and control NSG mice.

### The presence of human immune cells promotes dissemination of SARS-CoV-2 in HIS hACE2 KI NSG mice

hACE2 KI NSG HIS mice were generated as previously described (29) and subsequently stratified based on the inclusion (Hu/Hu) or omission (Hu) of human fetal thymus engraftment to support T cell reconstitution (Fig. 2a). Thymectomized hACE2 KI NSG mice that did not receive CD34+ HSCs or thymus transplants served as immunodeficient controls. As expected, both Hu/Hu and Hu mice had high percentages of reconstituted human CD45+ cells, detected via flow cytometry, in peripheral blood mononuclear cells, spleen, and digested lung tissue, while only Hu/Hu mice reconstituted human CD3+ T cells (Fig. 2b,c). Twelve to 16 weeks after transplantation, the mice were intranasally infected with either the WA1 strain of SARS-CoV-2, at a dosage of 5-10x10^3^ pfu (Fig. S1a-d) or with PBS as a control. Tissues were collected from mice at 7 and 14 days post-infection to model acute infection, and at 8 weeks post-infection to study the effects of long-term SARS-CoV-2 infection (Fig. 2a). Acutely infected mice did not demonstrate significant weight loss compared to uninfected controls, however in the long-term infection cohort there was significant weight loss in the infected group compared to uninfected controls (*p*-value = 0.03)(Fig. S1e). Thus, our model demonstrates gradual but sustained weight loss over a longer period of infection.

**Figure 2:**
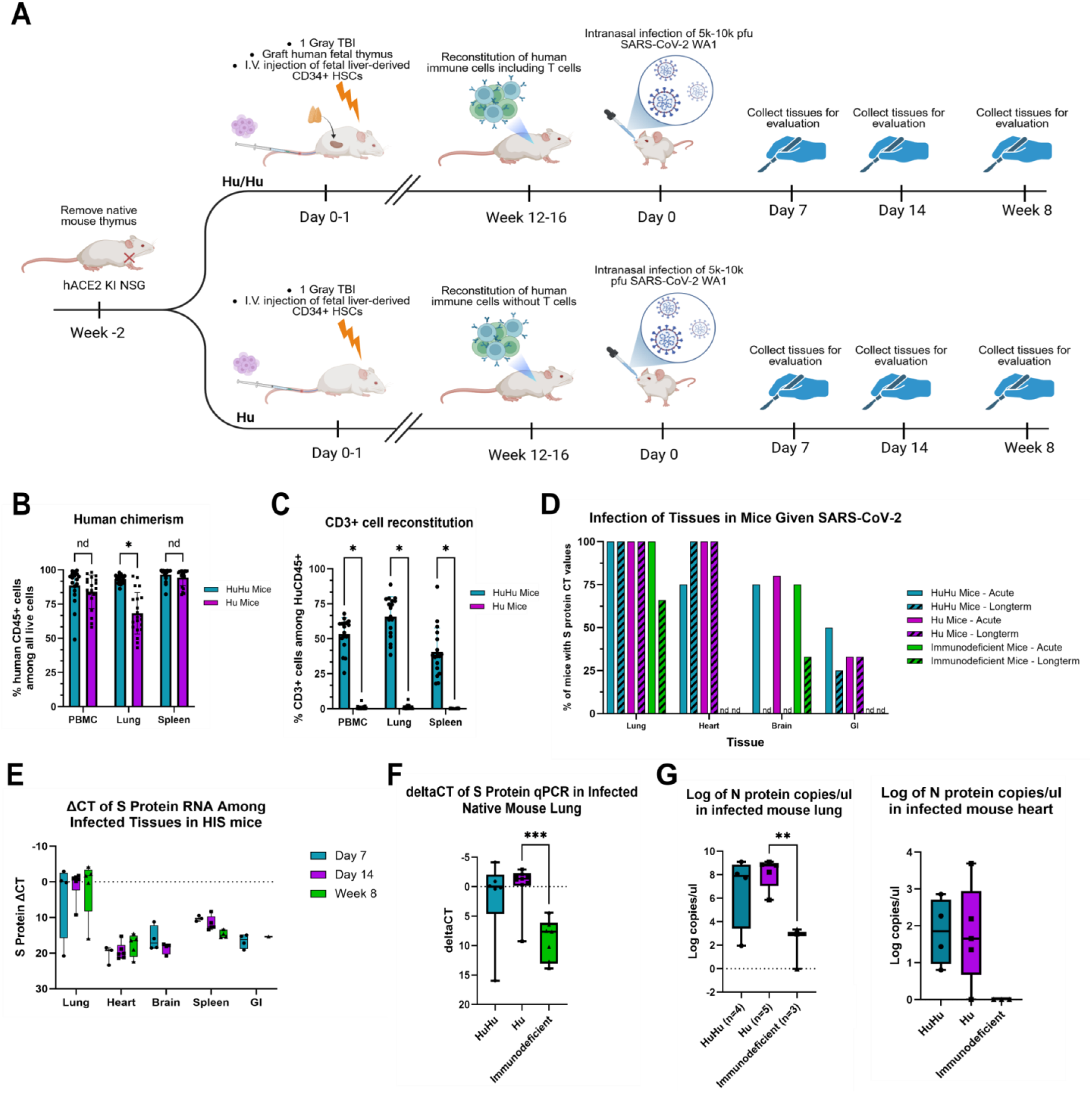
The presence of human immune cells promotes the dissemination of SARS-CoV-2 and increases viral levels in mouse lung. **a.** Schematic of hACE2 KI NSG HIS mouse infection study. A subset of mice also expressed an HLA-A2 transgene. Homozygous hACE2 KI NSG mice were thymectomized and reconstituted with either fetal liver CD34+ HSCs (2x10^5^ cells) alone or both HSCs and a human fetal thymus graft under the kidney capsule or left unreconstituted. After 3-4 months to allow human immune cell reconstitution, mice were intranasally infected with the WA1 strain of SARS-CoV-2 (5x10^3^ pfu) or mock infected with PBS. Tissues were harvested at 7, 14 days, and 8 weeks post-infection. **b,c.** Reconstitution of human CD45+ cells **(b)** and human CD3+ T cells **(c)** in PBMC, lung, and spleen of HIS hACE2 mice at 16-20 weeks post-transplantation, as determined by flow cytometry. Hu/Hu *n* = 19, Hu *n = 21.* All displayed *p*-values < 0.0001. **d.** Percentage of mice from each infected group demonstrating SARS-CoV-2 spike protein-specific qPCR CT values across multiple tissues. PBS control mice (not shown) did not exhibit spike protein-specific CT values. nd denotes tissues in which spike protein-specific CT values were not detected. **e.** ΔCT values of spike-protein specific qPCR normalized to mouse GAPDH across multiple tissues and both timepoints. Box and whisker plots show min to max values. Individual values for each mouse are shown. Day 7 *n* = 3-4, Day 14 *n* = 4-6, Week 8 *n* = 1-5. **f.** Levels of viral spike protein RNA detected in the native mouse lung for all infected mice across all timepoints. ΔCT values of SARS-CoV-2 spike protein-specific qPCR were determined by normalizing the spike protein values against mouse GAPDH. Welch’s ANOVA test *p*-value = 0.0014, with an adjusted *p*-value = 0.005 between Hu and immunodeficient groups. Individual values for each mouse are shown. Hu/Hu *n* = 6, Hu *n* = 9, Immunodeficient *n* = 7. g. Log copies per microliter in RNA isolated from lung (left) and heart (right) tissue from nucleocapsid protein-specific qPCR. Copy numbers determined using a standard curve from provided nucleocapsid standards. Box and whisker plots show min to max values with adjusted p-values determined by one-way ANOVA. Individual values for each mouse are shown. Lung: Hu/Hu *n* = 4, Hu *n* = 5, Immunodeficient *n* = 3. One-way ANOVA *p*-value = 0.0142. Heart: Hu/Hu *n* = 4, Hu *n* = 5, Immunodeficient *n* = 3.

Quantitative PCR for SARS-CoV-2 spike protein identified enhanced viral dissemination across most tissues in HIS mice compared to that in control hACE2 KI NSG mice lacking human immune cells, independent of infection duration (Fig. 2d). Notably, viral RNA was detected in hearts and intestines only of HIS mice and not in infected controls both in acute and long-term SARS-CoV-2 infection. Viral persistence signatures were observed across most tissues assayed, as spike protein transcript levels remained constant between timepoints (Fig. 2e). Of note, viral RNA was observed in the brain of all study groups across both acute timepoints but was absent at the 8-week timepoint in Hu/Hu and Hu mice, suggesting that clearance of the virus had occurred in the brain, but not in other tissues, in HIS mice over time.

The presence of human cells in reconstituted mice increased the overall viral RNA titers in the lungs compared to that in mice without human immune reconstitution in both acute and long-term infection (Fig. 2f, S1f). Nucleocapsid protein-specific RT-qPCR of the lung showed the same trend, with significantly increased viral titers detected in HIS mice compared to immunodeficient mice (Fig. 2g). Nucleocapsid protein RNA was also detected in plasma at 8 weeks post-infection in HIS mice only, further suggesting a role for human cells in the transportation of virus (Fig. S1g). Taken together, these data demonstrate prolonged viral persistence signatures across multiple organs in hACE2 KI HIS mice and suggest that human immune cell reconstitution influences viral dissemination.

### Infected HIS mice demonstrate COVID-19-related lung pathology and widespread viral protein in the lungs

Infected HIS mice exhibited acute lung injury reminiscent of COVID-19 patterns reported in human patients (21, 36–41), including fibrin formation, microthrombi and interstitial thickening. Similar pathology persisted at 8 weeks post-infection (Fig. 3a). In contrast, the lungs of uninfected HIS mice retained normal architecture.

**Figure 3:**
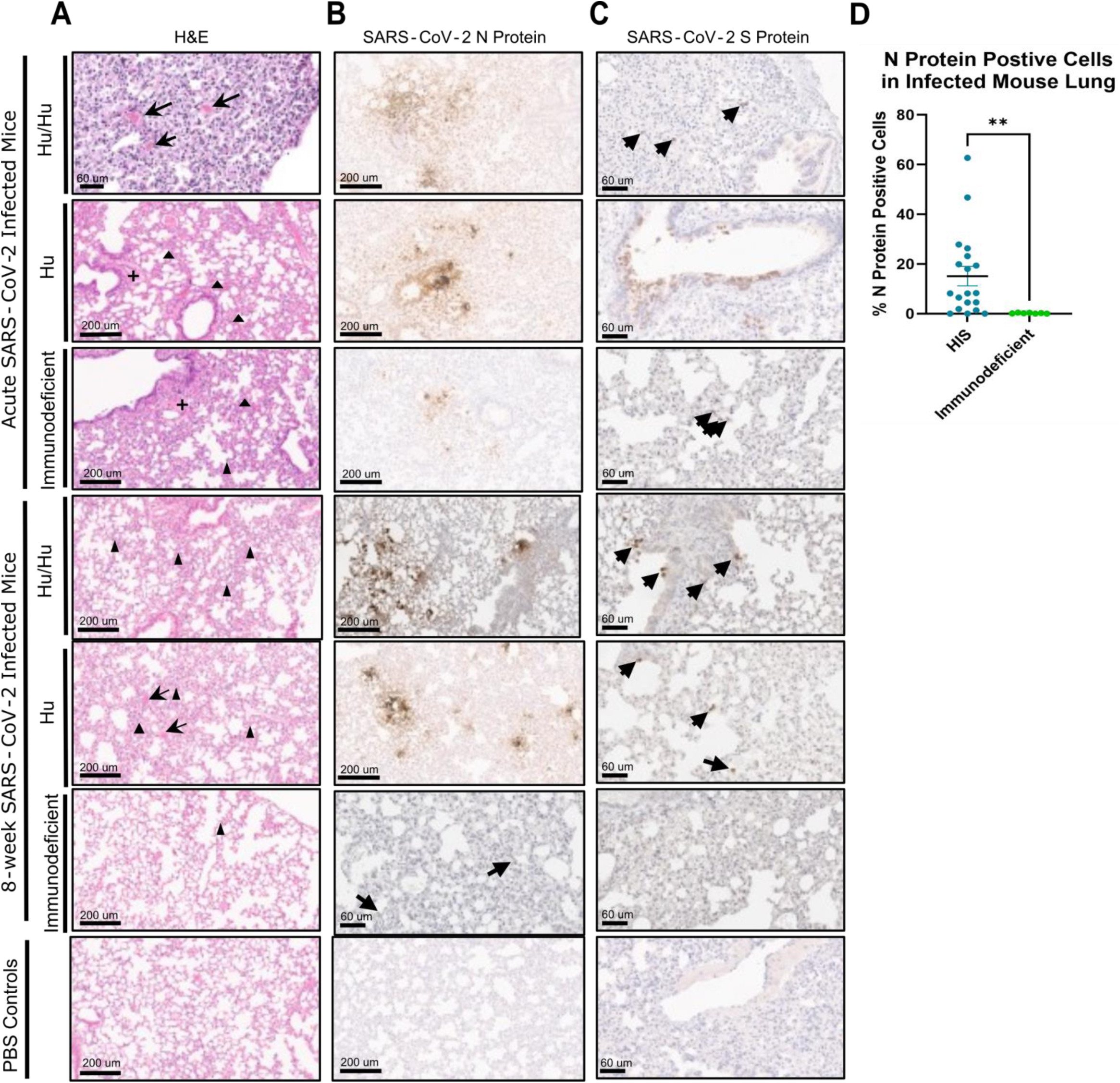
SARS-CoV-2 infected hACE2 KI NSG HIS mice demonstrate COVID-19-related lung pathology and widespread viral protein in mouse lung. **a.** Representative images of hematoxylin and eosin staining of formalin-fixed paraffin-embedded (FFPE) lung tissue from infected Hu/Hu (first and third rows), Hu (second and fourth rows), and immunodeficient mice (third and fifth rows) across both acute and long-term timepoints (indicated to the left of images), with an uninfected control mouse (bottom). Arrows indicate microthrombi, plus signs indicate fibrin formation, triangles indicate thickened interstitial regions. Images taken at 20X magnification except for acute Hu/Hu mouse, which is 40X. **b,c.** Representative images of SARS-CoV-2 nucleocapsid and spike protein staining of FFPE lung tissue. Images taken at 20X magnification (b) and 40X magnification (c). **d.** Percentage of SARS-CoV-2 nucleocapsid protein positive cells in infected HIS and immunodeficient mouse lung tissue calculated using HALO imaging software. Statistical significance was analyzed by Mann-Whitney test. *p-*value = 0.0027. HIS *n* = 19, Immunodeficient *n* = 7. Both acute and long-term timepoints combined.

Immunohistochemical (IHC) staining for SARS-CoV-2 spike and nucleocapsid proteins revealed widespread expression of viral proteins in the lungs in all infected groups, again persisting up to 8 weeks post-infection (Fig. 3b,c). Viral proteins were detected around airways and throughout the lung parenchyma. Nucleocapsid protein was present at higher levels than spike protein, consistent with human studies indicating that the nucleocapsid protein is a more sensitive indicator of infection (42, 43). These stains were both more prominent in HIS mice and HALO image analysis demonstrated a significantly higher percentage of nucleocapsid protein-positive cells in infected HIS mice compared to their immunodeficient counterparts (Fig. 3d, Fig. S2).

### Infected mouse lung demonstrates increased fibrotic remodeling compared to uninfected controls

To complement qualitative histopathological observations, we developed an unbiased computational pipeline to quantify fibrotic remodeling across 67 H&E-stained lung sections encompassing 307,118 tissue patches and 816,320 segmented cells (Fig. 4a). Patch-level features were extracted using the UNI2-h pathology foundation model, batch-corrected with ComBat to account for tissue processing differences, and clustered into 8 morphologically distinct tissue phenotypes by K-means. One cluster (C7) was enriched in infected slides and displayed hallmarks of fibrotic remodeling, including increased mesenchymal cellularity and elongated nuclear morphology (Fig. 4b). Whole-slide mapping of C7 patches revealed widespread fibrotic cluster distribution in infected lungs (C7 = 31.6-43.9% of tissue area) compared to near-absence in PBS controls (C7 = 0.0-0.2%), a pattern consistent across all three humanization backgrounds (Fig. S3a). We constructed a composite fibrosis score integrating three independent components: tissue-level fibrotic cluster proportion, mean nuclear eccentricity within fibrotic regions, and fibroblast proportion from cell-type classification. This composite score was significantly elevated in infected compared to PBS-treated lungs across multiple cohort definitions (Combined: p = 0.004, Mann-Whitney U; Acute: p = 0.008; Long-term: p = 0.024) (Fig. 4c, S3b). Bootstrap 95% confidence intervals for the infected-versus-control difference excluded zero in the combined and acute cohorts, and permutation testing confirmed significance (Combined p_empirical = 0.013; Acute p_empirical = 0.012).

**Figure 4:**
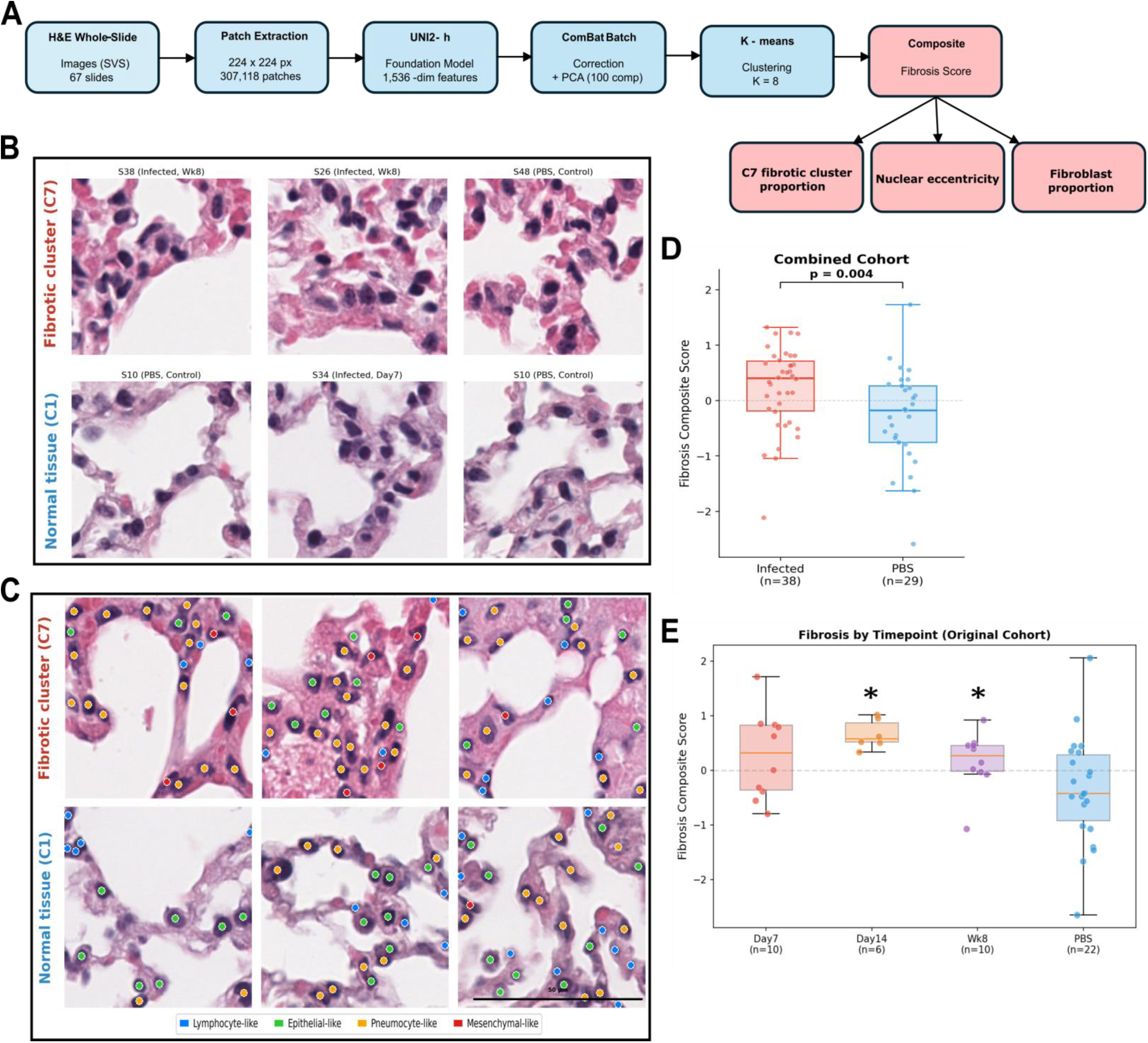
Infected mouse lung demonstrates increased fibrotic remodeling compared to uninfected controls. Computational histopathology quantifies fibrotic remodeling in SARS-CoV-2-infected lungs. **a.** Schematic of the computational pipeline: whole-slide H&E images (67 slides) were tiled into 307,118 patches, features extracted using the UNI2-h pathology foundation model, batch-corrected with ComBat, and clustered into 8 tissue morphology phenotypes. A composite fibrosis score was derived from three independent components. **b.** Representative patches from the fibrotic cluster (C7, top) and a normal tissue cluster (bottom). **c.** Cell segmentation with morphological phenotyping: individual nuclei in representative fibrotic (top) and normal (bottom) patches colored by morphological class — lymphocyte-like (blue), epithelial-like (green), pneumocyte-like (orange), and mesenchymal-like (red). Fibrotic tissue contains 5-15% mesenchymal-like cells compared to 0-3% in normal tissue. **d.** Composite fibrosis scores in infected versus PBS-treated lungs across all cohorts combined (p = 0.004, Mann-Whitney U test; n = 38 infected, 29 PBS). **e.** Temporal trajectory of fibrosis scores across Day 7, Day 14, and Week 8 post-infection, showing acute peak and persistent elevation. Holm-corrected *p-*value at Day 14 = 0.01, Week 8 = 0.022.

Individual score components were independently significant: fibroblast proportion (*p* = 0.007, Acute) and tissue fibrotic proportion (*p* = 0.005, Long-term 1), demonstrating that the composite captures converging histological signals rather than a single dominant feature (Fig. S3c). Stratification by humanization status revealed that fibrosis elevation was restricted to immune-reconstituted mice (Hu: p = 0.081; HuHu: p = 0.081), whereas immunodeficient controls showed no effect (ImDef: p = 0.345), consistent with an immune-mediated fibrotic process (Fig. S3d). Temporal analysis of fibrosis scores showed an acute peak at day 14 (mean = 0.659) compared to day 7 (mean = 0.277), with scores partially declining but remaining significantly elevated above PBS controls at week 8 (mean = 0.171), indicating persistent fibrotic remodeling beyond the acute phase (Fig. 4e). These quantitative findings corroborate our pathological observations and provide an unbiased, scalable framework for assessing lung fibrosis in preclinical SARS-CoV-2 models.

### Innate immune response is downregulated in mice with human T cells compared to those without T cells following infection

We evaluated transcriptional pathways in the lungs of acutely infected and uninfected Hu and Hu/Hu mice using the Nanostring platform and gene set enrichment analysis (GSEA). Pathway analysis, taking into account the overlap of differentially expressed genes in our samples with the full set of genes in each pathway, revealed the enrichment of multiple inflammasome signaling pathways, macrophage responses (MSP), NF-kβ, and IFN-γ responses in infected mice at day 7 post-infection. Leading genes in pathways with high enrichment scores included IL1b, CASP1, IFNGR2, CCL2, NFKB2, and JAK/STAT genes (Fig 5a, Table S1). At day 14 post-infection there was continued IFN-γ response and the emergence of innate inflammation-centered pathways. Anti-viral inflammatory response pathways such as TLR signaling, NF-kβ activation and macrophage stimulation were enriched in infected lung samples. The leading genes in the top-most enriched pathways included toll-like receptor genes (TLR1/2/3/4/6/8), MYD88, TNFSF10, JAK2, IFN-γ, and NFKB2 (Fig 5b, Table S2).

**Figure 5:**
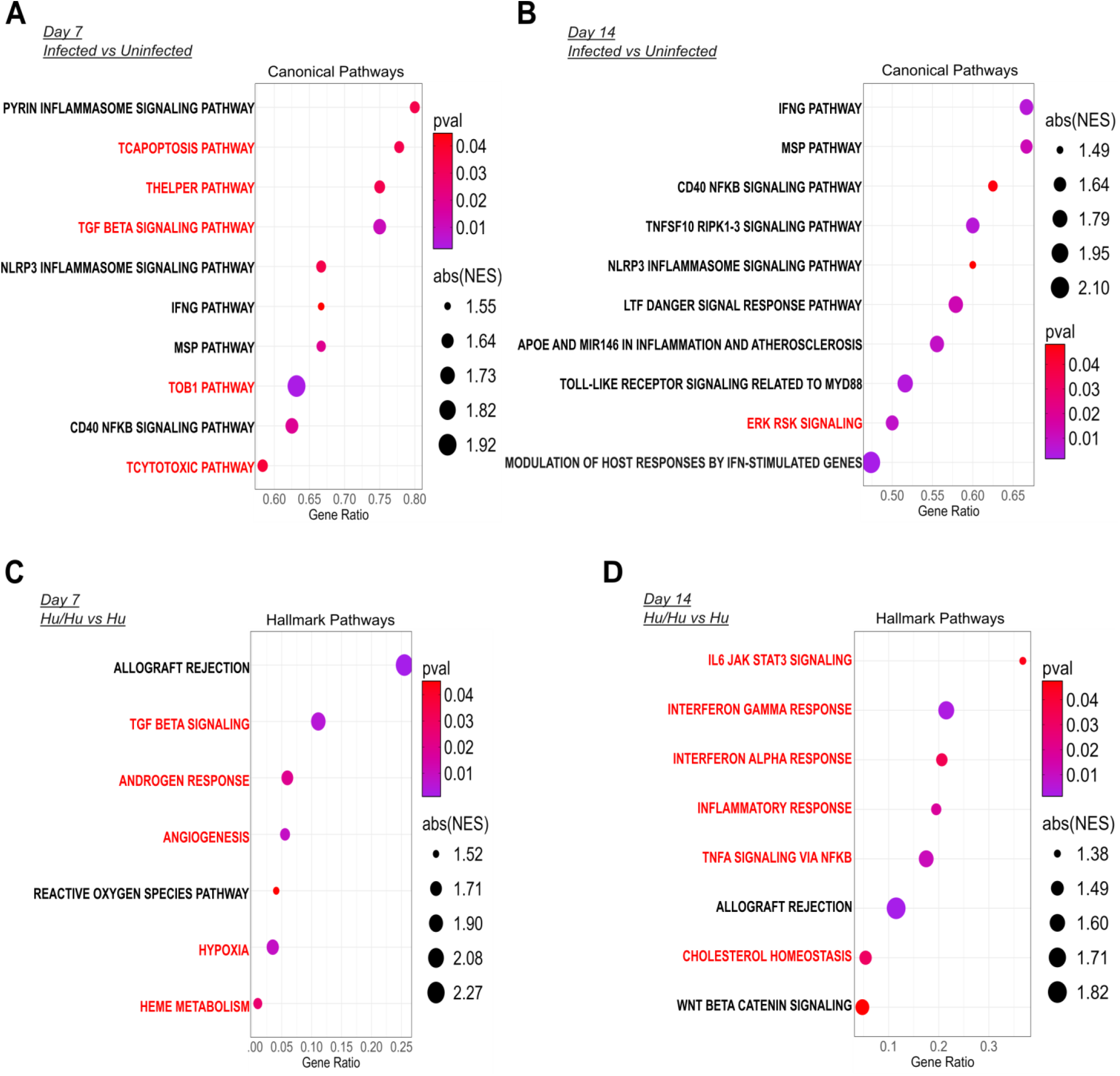
Infected mice without T cell reconstitution demonstrate a more pronounced IFN response in the lungs than infected mice with T cell reconstitution. Human T cells delay innate antiviral responses in SARS-CoV-2 infection. **a,b.** Dot plot showing the most highly enriched pathways utilizing the canonical pathway geneset from MSigDB as identified by GSEA between infected and uninfected HIS mouse lung. Data includes all Hu and Hu/Hu mice at 7 (infected *n* = 10, uninfected *n* = 12)(a) and 14 days (infected *n* = 12, uninfected *n* = 12) (b) post-infection. Red pathway text denotes downregulation of the pathway while black pathway text denotes upregulation of the pathway in infected mice. The x-axis denotes the gene ratio of enriched genes in the pathway versus total genes in the pathway. Size of dot denotes absolute normalized enrichment score with color of dot denoting the significance. **c,d.** Dot plot showing the most highly enriched pathways utilizing the human hallmark pathway geneset from MSigDB as identified by GSEA between infected Hu/Hu and Hu mouse lung at day 7- (Hu/Hu *n* = 5, Hu *n* = 5) (c) and 14-days (Hu/Hu *n* = 5, Hu *n* = 5) (d) post-infection.

To address the potential cross-reactivity of the Nanostring probes between human and murine genes, we evaluated their capture of immunodeficient mouse RNA in an infected unreconstituted hACE2 KI NSG mouse (Fig. S4). 56.4% of Nanostring probes (436 of 773) had minimal murine cross-hybridization (<10% of signal from human samples), while an additional 25.9% showed moderate cross-reactivity (10-50%). Critically, 87.5% of IFN/JAK-STAT pathway genes (42 of 48) showed less than 10% cross-reactivity, with key interferon-stimulated genes (MX1, IFIT1-3, ISG15, OAS1-3, GBP1-5, CXCL10/11) all below 5% cross-reactivity. Only two IFN-related genes (DDX58 and IFNB1) exceeded 50% cross-reactivity. These results suggest that the majority of the response indicated by the Nanostring data is derived from human cells.

We then compared immune responses between infected Hu and Hu/Hu mice, to investigate how the presence or absence of T cells may affect the response to acute infection. Differential gene expression (DGE) analysis identified 122 differentially expressed genes (DEGs) between these groups at day 7 and 55 at day 14, with the majority relating to the presence of T cells in Hu/Hu mice (Fig S5). These T cell-related DEGs included TCR signaling genes (ZAP70, TRAT1), cytotoxic genes (GZMA, KLRB1), costimulatory genes (CD40LG, CD28, ICOS, LCK, ITK), T cell differentiation markers (TCF7 and LEF1), and exhaustion markers (CTLA4 and LAG3). Pathway analysis revealed an enriched T cell response in infected Hu/Hu mice at both timepoints and downregulation of IFN responses on day 14 compared to infected Hu mice (Fig 5c,d). TGFb signaling was enriched in the Hu population at day 7, shifting into an IFN-based response by day 14 (Fig 5c,d). Leading genes in the allograft rejection pathway at both days 7 and 14 included CD3D, CD3E, TRAT1, LCK, ITK, CD28 and CD3G, reflecting the presence of reconstituted T cells in Hu/Hu mice and including many of the T cell-related DEGs mentioned previously (Table S3-4). Leading genes involved in pathways more enriched in Hu mice than Hu/Hu mice included THBS1 and SMAD3 in the TGFb signaling pathway at day 7, shifting to interferon-inducible genes (IFIT2/3, IFI44, IFI27) and innate response genes such as RIPK2, CXCL3/8, IL6, ISG15, MYD88 and TLR2 by day 14. Additionally, IFN-γ was not among the leading genes in the IFN gamma response pathway, with the enrichment instead influenced by IFI and innate anti-viral response genes. These differences elucidate the contrast of adaptive versus innate antiviral response in mice with and without T cell reconstitution, with Hu mice demonstrating a larger innate antiviral response including genes prominently implicated in COVID-19 pathogenesis (44). Despite the presence of T cell-defining genes, markers of TCR signaling, and evidence of a cytotoxic response through the presence of GZMA, Hu/Hu mice did not clear infection. However, while Hu mice demonstrated a strong anti-viral and pro-inflammatory response, with upregulation of IL-6 and IFI genes, this response was also insufficient to clear acute infection.

### Myeloid expansion and lymphocyte depletion is observed in the lungs of infected mice

Single nucleus RNAseq/TCRseq was performed on nuclei from lungs of infected and uninfected mice at 8 weeks post-infection. Analysis of human nuclei revealed distinct populations of T cells, B cells, monocytes, macrophages, and dendritic cells (Fig. 6a, Fig. S6a). Compositional analysis of these populations revealed a relative expansion of myeloid cells and depletion of lymphoid cells in infected mice compared to uninfected controls, mirroring what has been seen in COVID-19 patient BAL studies (Fig. 6b-c, S6b) (45, 46). Bootstrap downsampling of these populations confirmed a 29.6% increase in the monocyte population in infected samples compared to uninfected samples and a decrease in T cell and B lineage cell (including plasma cells) populations by 20.1% and 17% respectively. These data were corroborated via flow cytometry, which revealed significant decreases in HuCD45+CD3+ cells and CD4+ T cells in infected compared to uninfected Hu/Hu mouse lung (Fig. S6c) and spleen (Fig. S6d) samples at 8 weeks post-infection.

**Figure 6:**
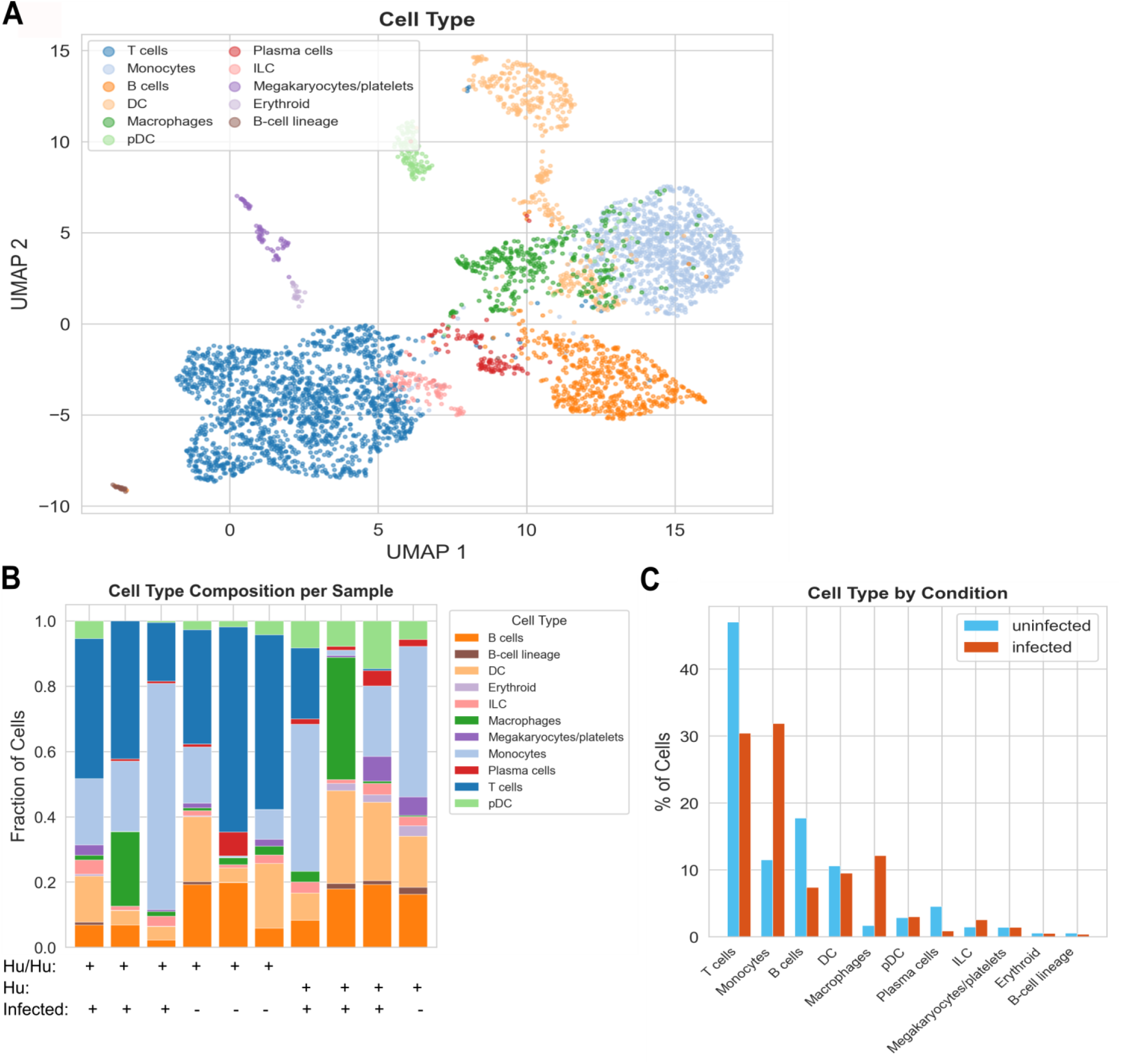
Myeloid expansion and lymphoid depletion is observed in infected mouse lung. **a.** UMAP of 4,401 human immune lung cells colored by cell type annotation. Cell annotations were generated by CellTypist using the Immune All High reference. **b.** Cell type composition comparing infected and uninfected Hu/Hu and Hu mice, displayed as stacked barplots. **c.** Breakdown of the percentage of cell populations in infected (red) and uninfected (blue) Hu/Hu mice, displayed as a bar plot.

### Lung-infiltrating human CD4+ T cells demonstrate a marked shift towards cytotoxic programming after long-term infection

To gain further insight into T cell responses to infection, we conducted differential expression analysis on CD4+ T cells obtained from snRNA-seq of Hu/Hu mouse lung at 8 weeks post-infection. Strongly upregulated DEGs in the CD4+ T cell compartment of infected mice included PARP14, ITGA4, PCNX1, and DOCK8- genes involved in IFN I responses, T cell activation, and leukocyte trafficking, suggesting an immune response to infection and trafficking into infected lung tissue (Fig. 7a). LEF1, BCL2, and TOX were downregulated in infected mice, reflecting a loss of naïve programming and decreased T follicular helper cell differentiation along with increased T cell apoptosis (47, 48). Transcription factor activity driving the CD4+ T cell response included SOX5, which is induced by IL-6 and the STAT3 pathway and drives pro-inflammatory Th17 differentiation (Fig. 7b) (49). Sox9, TCF7, and GATA3 were significantly downregulated in infected animals, confirming loss of naïve T cell identity and decreased T cell quiescence (47, 50, 51). This marks a significant shift from the acute data, where TCF7 was upregulated in infected Hu/Hu lung samples at both days 7 and 14. Pathway analysis of CD4+ T cells in infected and uninfected Hu/Hu mice CD4+ T demonstrated IFN pathway and JAK/STAT pathway activation (Fig. 7c), indicating an active immune response and showing a marked shift from the response observed in acutely infected Hu/Hu mice, where an IFN response was not evident (Fig. 5b,c). PARP14, MX1, IFI44L, SP100 and EIF2AK2 were the top upregulated interferon-stimulated genes in 8-week infected samples, many of which have been commonly identified DEGs in respiratory infections (52).

**Figure 7:**
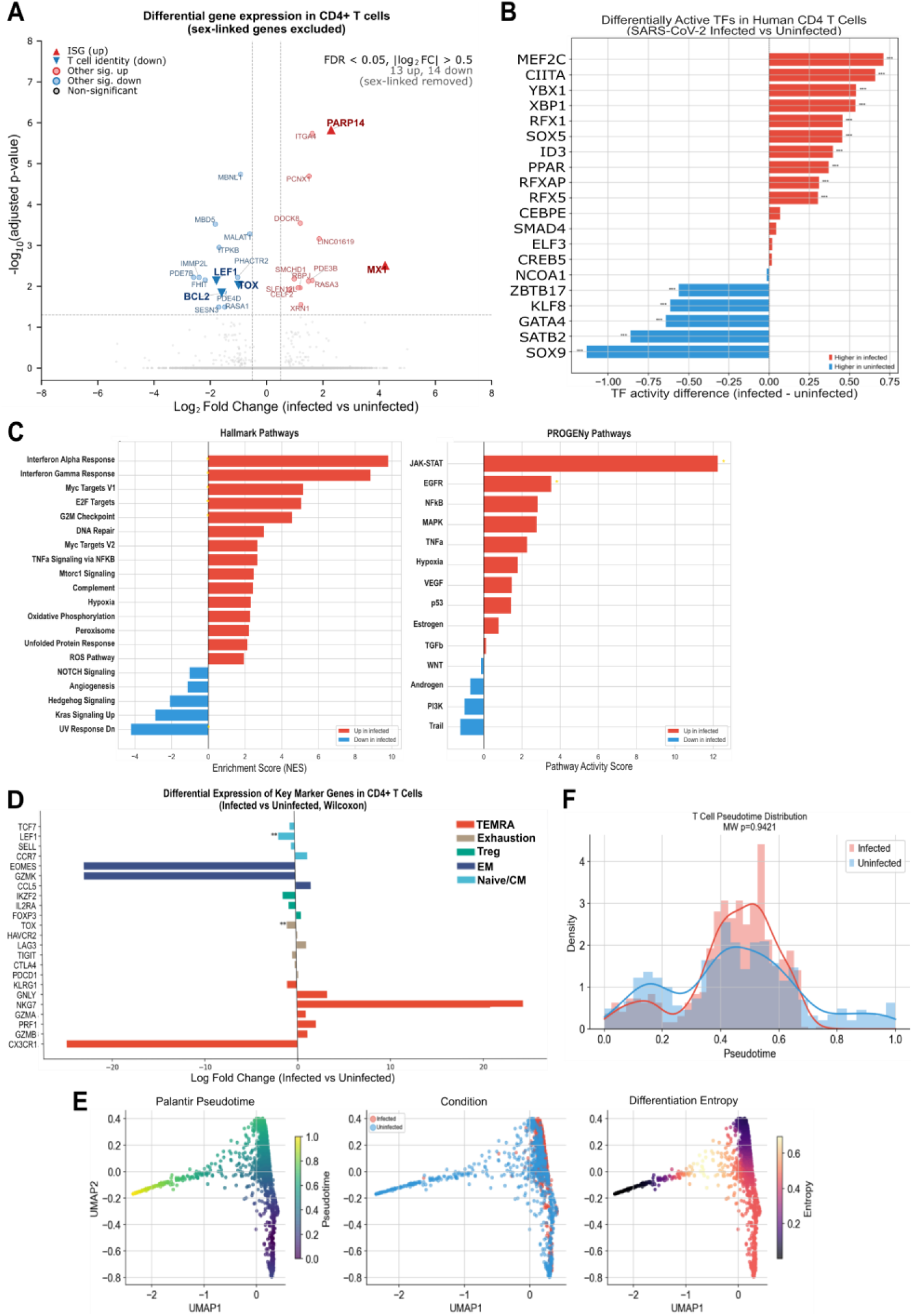
CD4 T cells in infected mice demonstrate IFN and JAK/STAT pathway activation and a shift towards cytotoxic program activation compared to uninfected mice. **a.** Volcano plot of the 14,260 autosomal genes tested in CD4⁺ T cells (n = 271 cells) using the Wilcoxon rank-sum test implemented in ‘scanpy.tl.rank\_genes\_groups’ with Benjamini-Hochberg correction; X- and Y-chromosome genes were identified by Biomart query against the Ensembl GRCh38 annotation and excluded (n = 439 removed). Dashed grey lines mark the significance thresholds (horizontal: adjusted p = 0.05; vertical: |log₂ fold-change| = 0.5). Each gene passing both thresholds is labeled: interferon-stimulated genes are highlighted as large red upward triangles (PARP14, log₂FC = +2.29, p_adj = 1.5 × 10⁻⁶; MX1, log₂FC = +4.21, p_adj = 3.0 × 10⁻³), and canonical T-cell identity or survival markers are highlighted as large blue downward triangles (LEF1, log₂FC = −1.78, p_adj = 7.4 × 10⁻³; BCL2, log₂FC = −1.59, p_adj = 1.5 × 10⁻²; TOX, log₂FC = −0.99, p_adj = 9.5 × 10⁻³). Other significantly up-regulated genes (n = 11, muted red) and down-regulated genes (n = 11, muted blue) are shown with smaller markers and labeled in a correspondingly muted font. **b.** Top differentially active TFs in human CD4 T cells as determined by decoupler ULM with CollecTRI regulons. **c.** Enrichment scores denoting the most highly enriched pathways in CD4 T cells in infected Hu/Hu mouse lung at 8 weeks post-infection, utilizing the Hallmark (left) and PROGENy (right) databases at the single nuclei level. IFN-alpha response *p* < 1e−19, IFN-gamma response *p* < 1e−15, JAK-STAT signaling *p* < 1e−31. **d.** Differential expression of key CD4 T cell markers and cytotoxic effector genes comparing infected and uninfected Hu/Hu mouse lung. Overall subset scores computed using scanpy.tl.score_genes for each subset using the denoted key marker genes. P-values calculated by Mann-Whitney U test (two-sided). Naïve: p<0.001, down. Central memory: p=0.002, down. **p-value < 0.01. **e.** Human T cell trajectory on diffusion components. **f.** Pseudotime density distributions by condition. Infected cells shift toward higher pseudotime values (KS *p*< 0.0001).

To further investigate these differences in the context of distinct T cell populations, we used the differential expression of key marker gene sets to look at CD4+ subsets, which revealed a significant depletion of the naïve (p<0.001) and central memory (p=0.002) CD4+ T cell compartments, influenced by the significant decrease in LEF1 expression (padj = 0.007) alongside TCF7. Through this T cell subset analysis, a “TEMRA-high” population was transcriptionally identified using mature T cell markers (CX3CR1, GZMB, PRF1, GZMA, KLRG1, NKG7, GNLY, FGFBP2). Infected samples exhibited upregulation of the majority of these genes, which characterize a distinct CD4+ CTL population that has been observed in responses to injury and infection (53, 54). An increase in the TEMRA population was also confirmed via flow cytometry in infected Hu/Hu mice at 8 weeks post-infection, validating this population both transcriptionally and via surface markers (Fig. S6e). We used pseudotime ordering, choosing root cells based on the highest naïve signature (CCR7, LEF1, TCF7, and SELL), to view the trajectory of this CD4+ T cell compartment (Fig 7e-f). This analysis confirmed a significant shift in infected CD4+ T cells (KS p<0.0001) towards higher, more effector-like pseudotime values.

### CD8+ TEMRA and TRM cells are increased in long-term infected Hu/Hu mice

Analysis of the CD8+ T cell compartment revealed similar trends to those observed in the CD4+ T cell compartment. Significant decreases in naïve (p=0.003) and central memory (p=6.8x10^-4^) CD8+ T cells appeared in infected Hu/Hu mice at 8 weeks post-infection (Fig. S7a). However, increases in the TEMRA (p=0.015) and tissue resident memory (TRM) cell compartment (p=0.004) were also observed, the latter suggesting formation of tissue-resident memory in the lung in response to infection. Pathway analysis revealed enrichment of IFN-response pathways (p<0.05) and JAK-STAT pathways (Fig. S7b). Thus, both CD4+ and CD8+ T cell compartments demonstrate a shift towards mature, effector transcriptional programming with upregulation of pro-inflammatory pathways. Additionally, flow cytometry analysis of CD8+ T cells in infected lung and spleen showed upregulation of PD1 at 8 weeks post-infection (Fig. S7d).

### SARS-CoV-2-specific T cell responses in infected Hu/Hu mice

To evaluate whether infected Hu/Hu mice developed SARS-CoV-2-specific T cell responses, we first used splenocytes collected from acutely infected mice to perform an ELISpot assay against SARS-CoV-2 peptide pools. At day 7 post-infection we detected responses to a range of SARS-CoV-2 spike protein, nucleocapsid protein, envelope protein and membrane protein peptides (Fig. 8a). These SARS-CoV-2-specific T cell responses persisted in infected Hu/Hu mice at 8 weeks post-infection, demonstrating a long-lasting antigen-specific response (Fig. 8b).

**Figure 8:**
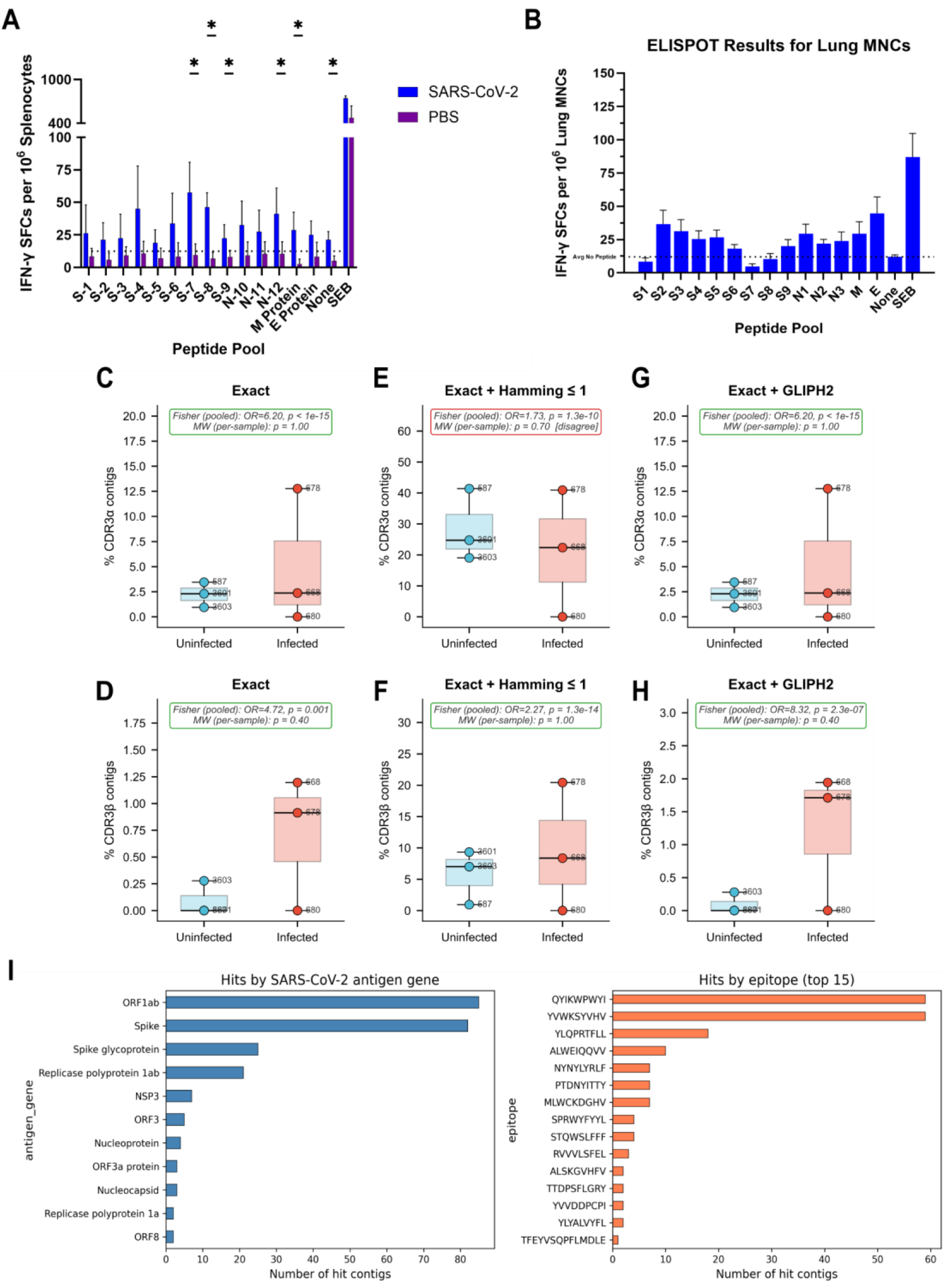
Infected Hu/Hu mice demonstrate SARS-CoV-2-specific TCR enrichment. **a.** T cell specific responses against SARS-CoV-2 spike, nucleocapsid, membrane and envelope proteins, determined by human IFN-γ ELISpot assay at 7 days post-infection in splenic T cells. Peptide pools indicate 17-mer overlapping peptides. Means with standard deviation are plotted. Unpaired t-tests were performed between groups for each peptide pool, with asterisks denoting significant differences with p<0.05. Dotted line denotes average number of spots without stimulation. Responses are considered positive above this line. SARS-CoV-2 *n* = 4, PBS *n* = 10. **b.** T cell specific responses against SARS-CoV-2 peptide pools in isolated lung mononuclear cells of infected Hu/Hu mice at 8 weeks post-infection. Peptide pools indicate 17-mer overlapping peptides. Error bars indicate mean ± SEM. Dotted line denotes average number of spots without stimulation. Responses are considered positive above this line. **c-h.** T-cell receptor α (CDR3α) and β (CDR3β) chain contigs (TCR sequences) assembled by TRUST4 from 10x VDJ data of six Hu/Hu mouse lung samples (n = 3 infected, 3 uninfected) were matched against a combined SARS-CoV-2 reference built from VDJdb, McPAS-TCR, and IEDB using three successively more permissive tiers: **(c,d)** exact CDR3 amino-acid identity, **(e,f)** Hamming distance ≤ 1 (same length, one substitution), and **(g,h)** GLIPH2 motif sharing (a sample CDR3 shares a short motif enriched in SARS-CoV-2 reference CDR3s relative to non-SARS human CDR3s; see Methods). Bars are colored by condition: blue = uninfected, red = infected. Box plots show per-sample rates (median, IQR, whiskers); individual sample IDs are annotated. Each panel reports two statistical tests: Fisher’s exact test on pooled contigs (the contig-level test; odds ratio and two-sided p-value) and Mann-Whitney U on per-sample rates (the sample-level test; n = 3 vs 3, minimum achievable two-sided p ≈ 0.1). Panel borders are colored green where both tests agree on direction and red where they disagree. **i.** SARS-CoV-2-specific antigen genes and epitopes detected by TCR contigs from 8-week infected Hu/Hu mice. Number of contigs based on antigen target (left) and known SARS-CoV-2-specific epitope sequence (right).

We then investigated whether T cell receptor (TCR) sequences isolated from digested lung tissue during paired TCR and single nuclei RNA-sequencing from mice at 8 weeks post-infection matched known SARS-specific TCR sequences. We compared 3,891 TCRβ contigs (TCR sequences) sequenced from 6 Hu/Hu mice (infected *n* = 3, uninfected *n* =3) to sequences recognizing known epitopes isolated from human SARS-CoV-2 patients (55). TCRs exactly matching known SARS-CoV-2 TCRs were significantly enriched in both CDR3α (p<0.001) and CDR3β (p=0.001) sequences from infected mice (Fig. 8c,d). SARS-CoV-2-specific CDR3β sequences, but not CDR3α sequences, were also further enriched in infected mice by a Hamming distance of one or fewer amino acid substitutions (*p*<0.001) (Fig. 8e,f). While no GLIPH2 motifs were detected in CDR3α contigs, CDR3β contigs in infected mice demonstrated enrichment for amino acid motifs present in human TCR sequences isolated from SARS-CoV-2 patients (p<0.001) (Fig. 8g,h). The top clonally expanded SARS-specific TCRs isolated from these mice targeted ORF1ab and SARS-CoV-2 spike glycoprotein epitopes, consistent with those predominantly detected in human patients post-infection (Fig. 8i)(55). Thus, hACE2 KI Hu/Hu mice demonstrate long-lasting antigen-specific TCR responses to SARS-CoV-2.

### Transcriptional analysis of mouse lung nuclei reveals a chemokine-driven coordinated tissue response to infection

To investigate long-term transcriptional responses in the lung tissue of infected mice, we performed single nucleus RNA-sequencing on dissociated lung tissue from a subset of mice (infected Hu/Hu n = 3, uninfected Hu/Hu n = 3, infected Hu n = 3, uninfected Hu n = 1, infected immunodeficient n = 3, and uninfected immunodeficient n = 2) at 8 weeks post-infection. Each nucleus was identified as being of human or mouse origin if >80% of barcoded transcripts mapped to one species. Downstream analysis of each species was completed separately.

Analysis of the murine nuclei revealed major cell populations such as AT1 and AT2 alveolar epithelial cells, ciliated cells, secretory airway epithelium, capillary endothelial cells (CAP1/CAP2), and alveolar macrophages (Fig. 9a, S8a). Compositional analysis of these populations revealed a significant decrease in ciliated cells in infected mice compared to uninfected controls (p=0.011), along with a substantial decrease in the secretory cell population (p=0.057) (Fig. 9b). We also observed a decrease in cell-cell communication between ciliated cells and other cell populations such as AT1-AT2 cells and alveolar macrophages using receptor-ligand analysis, further confirming the loss of ciliated cells in infected mice (Fig. S8b).

**Figure 9:**
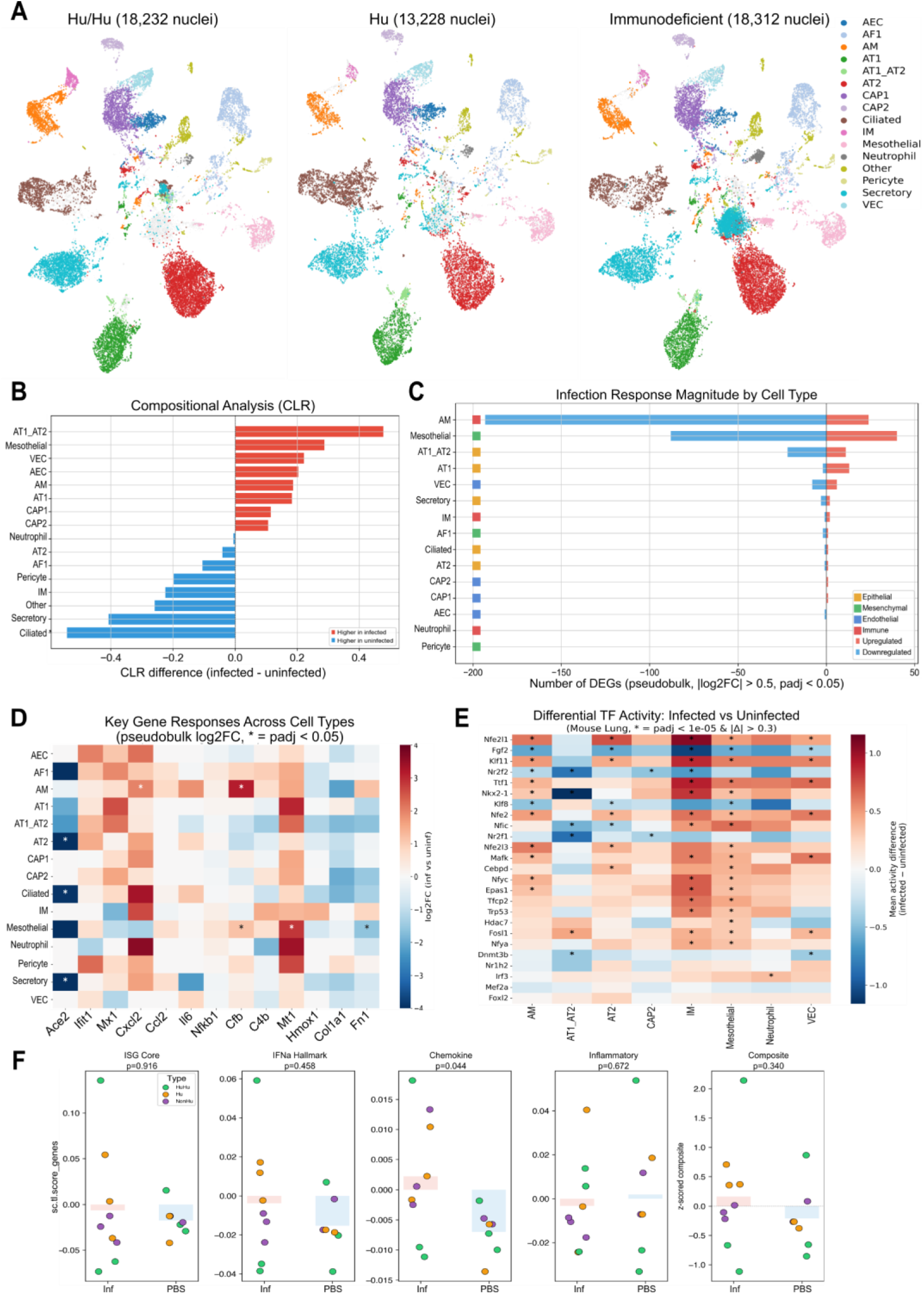
Infected mouse tissue response is dominated by macrophage activation. **a.** UMAP of 49,772 mouse lung cells split by humanization type (HuHu, Hu, Immunodeficient). Cell type annotations were generated using CellTypist with two reference atlases (LungMAP MouseLung CellRef v1.1 and Mm-Flu-timecourse). **b.** Centered log-ratio transformation of mouse lung cell compositional analysis. Ciliated cell CLR p=0.011, Ciliated cell scCODA inclusion probability = 0.91. **c.** Number of differentially expressed genes per cell type (pseudobulk, |log2FC| > 0.5, *padj* < 0.05). Red = upregulated, blue = downregulated. Colored by lineage. **d.** Log2FC of biologically important genes across cell types. Stars indicate *padj* < 0.05. **e.** Top differentially active TFs per mouse lung cell type (infected vs uninfected). Color = mean activity difference; * = *padj* < 1e-5, with a mean activity difference > 0.3. **f.** A priori infection response scores using published, dataset-independent gene sets. *p-*values displayed at the top of each graph.

To further investigate the transcriptional response to infection, we performed pseudobulk differential expression analysis and identified alveolar macrophages as the primary responder to infection, with 217 genes differentially expressed between infected and uninfected mice (Fig. 9c). Within alveolar macrophages, complement (Cfb), chemokines (CXCL2), and stress response genes (Mt1) were upregulated in response to infection, with significant transcriptional upregulation of NFkB1 (padj < 10^-22^) and PPARG (padj < 10^-21^), indicating a strong murine inflammatory macrophage response to infection (Fig. 9d,e). Ace2 was also downregulated across epithelial cells in infected mice including AT2 cells, ciliated cells, and secretory cells (Fig. 9d), suggesting infection of these cells, as a similar finding has been observed across multiple coronavirus infections (56–58).

To identify the specific genes driving response to infection, we performed a priori gene set scoring using four published gene sets involving ISG signatures, IFN-α response, chemokine response, and inflammatory response (59, 60). Only the chemokine response score significantly separated infected and uninfected samples (p=0.044), demonstrating that the infection response in the mouse lung is a chemokine-dominant instead of an IFN-dominant response (Fig. 9f). This might reflect a failure of murine cells to respond to human immune cell-derived cytokines due to differences in cytokine receptors between mice and humans(22). Spearman correlation of 55 differentially expressed genes present in two or more cell types revealed that the infection response was coordinated across multiple cell types, as all pairwise correlations were strongly positive (Spearman p=0.56-0.86) (Fig S8c). Stratifying the response by humanization status, we found that even immunodeficient mice showed a response to infection in alveolar macrophages (Fig. S8d). However, Hu/Hu and Hu mice had a broader response ranging across cellular compartments and a larger number of differentially expressed genes overall, indicating the effect of human immune cells on the murine response to infection at the tissue level.

### Infected HIS mice show multi-organ response to infection

Nanostring analysis revealed the presence of viral RNA in distal organs at both acute and long-term timepoints and GSEA using the canonical pathways database also revealed broad enrichment of anti-viral response pathways in heart tissue of infected mice compared to PBS-treated controls (Fig. 10a) (61). Acutely infected mice were enriched for innate anti-viral and IFN-response pathways, similar to observations in the infected lung. The leading genes in pathways with high enrichment scores involved JAK-STAT signaling genes, along with IFN-γ, IFNAR1, IL6, TLR genes, MYD88 and NFKB1 (Table S5). TBK1 and OAS3 were also leading genes in the Type I IFN induction pathway, and both have been implicated in COVID-19 pathology (62, 63). By the 8-week timepoint, IFN response pathways were absent (Fig. 10b). However, there was enrichment for both an inflammasome signaling pathway involving CASP1, IL1B, and NLRC4 and macrophage markers involving CD68 and CD163 (Table S6), suggesting ongoing innate immune activation in response to virus, as has been observed in PBMC samples in other studies (28, 64). There was also continued upregulation of NFKB1, IL6 and TNF in the heart at 8 weeks post-infection, providing evidence for an ongoing immune response.

**Fig 10.**
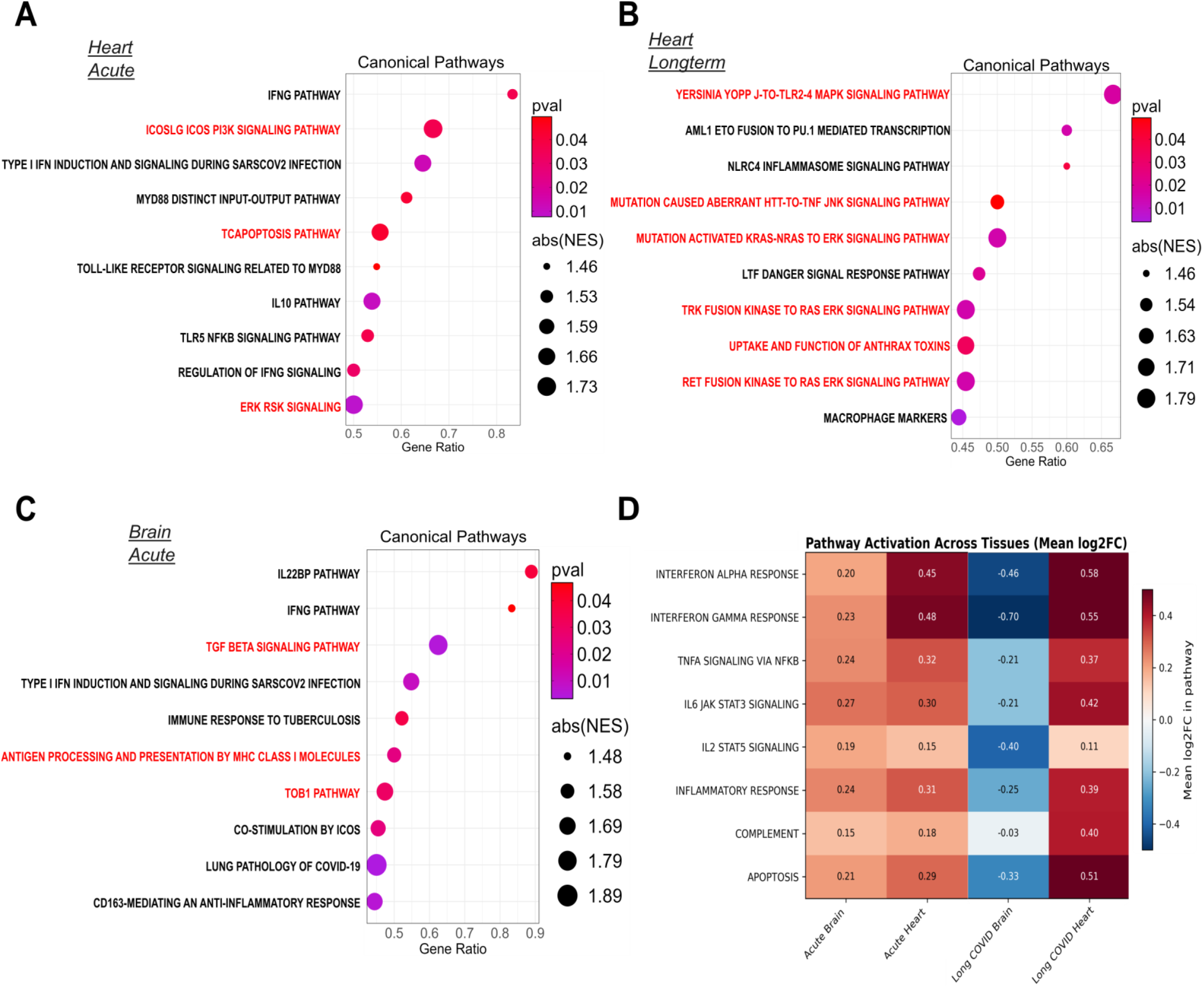
Infected HIS mice demonstrate evidence of multi-organ response to infection. Dot plots showing the most highly enriched canonical pathways as identified by GSEA in infected and uninfected mouse heart at acute (infected *n* = 3, uninfected *n* = 3) (**a**) and longterm (infected *n* = 6, uninfected *n* = 3) (**b**) timepoints, combining both Hu and Hu/Hu mice. **c.** Dot plot showing the most highly enriched canonical pathways as identified by GSEA in infected and uninfected mouse brain (infected *n* = 6, uninfected *n* = 5), combining all Hu and Hu/Hu mice. Red pathway text denotes downregulation of the pathway while black pathway text denotes upregulation of the pathway in infected mice. The x-axis denotes the gene ratio of enriched genes in the pathway versus total genes in the pathway. Size of dot denotes absolute normalized enrichment score with color of dot denoting the significance. **d.** Competitive GSEA across all heart and brain datasets using the Hallmark pathway dataset. Color = mean log2FC in pathway. IFN-gamma (padj = 0.007), IFN-alpha (padj = 0.010), and JAK-STAT (padj = 0.049) pathways were significantly enriched in acute heart tissue.

In homogenized brain tissue, acutely infected mice demonstrated enrichment of immune pathways such as the IFN-γ pathway, Type I IFN induction, and multiple disease response pathways (Fig.10c). Leading genes in these enriched immune response pathways included members of the JAK-STAT signaling pathway, TNF, IFN-γ, MX1 and IFIH1 (Table S7). OAS1, an interferon-induced enzyme that is produced in response to viral dsRNA, was one of the leading genes in the type I IFN pathway. OAS1 loss-of-function polymorphisms have been linked to both more severe COVID-19 outcomes, through the inhibition of anti-viral responses, and Alzheimer’s disease (65, 66).

Additionally, acutely infected brain samples also demonstrated enrichment of macrophage-mediated inflammatory responses involving CD163, IL6, and IL10. We did not detect viral RNA in the brain at 8 weeks post-infection and did not see evidence of overt immune responses in brains of infected mice through GSEA of Nanostring data. Consistently, competitive gene set enrichment analysis utilizing the Hallmark pathway dataset from MsigDB showed enrichment of IFN, JAK/STAT signaling, and complement pathways in acutely infected heart and brain samples and long-term infected heart, but not brain, samples (Fig. 10d). Overall, these data suggest a prominent immune response to SARS-CoV-2 infection across a broad range of tissues in infected HIS mice.

## Discussion

Here we describe a new human immune system model with prolonged viral persistence signatures across multiple organs. Due to its immunodeficient NSG background and physiologic expression of hACE, respectively, the hACE2 KI NSG HIS mouse model permits the assessment of human immune responses to SARS-CoV-2 in multiple tissues. We demonstrate a role for human immune cells in disseminating the virus, along with development of an ineffective T cell response that fails to clear virus while suppressing innate anti-viral IFN responses to infection. Our results replicate many of the features of COVID-19 and PASC, including lung pathology (41, 67, 68), viral dissemination and persistence in multiple tissues (69–72) and a long-lasting inflammatory response in these tissues (7, 73, 74). This unique small animal model provides an opportunity to better understand the role of virus and human immune responses in driving COVID-19 and PASC.

The wide distribution of SARS-CoV-2 RNA enabled by physiologic hACE2 expression in the knock-in mice resembles the distribution of viral antigen seen in decedent patients (75). This infection pattern was associated with widespread antiviral immune responses in heart and brain tissue. Remarkably, the presence of human cells increased viral loads in the lungs and extrapulmonary tissues compared to non-reconstituted mice. Similar viral loads detected in Hu and Hu/Hu animals indicate that T cells are not required for viral dissemination or replication. Acutely infected HIS mice demonstrated enrichment of inflammasome activation pathways, including upregulation of CASP genes and NLRP3, in human macrophages, consistent with *in vitro* and *in vivo* studies in HIS mice receiving inhaled AAV-hACE2 and in patients (28, 64, 76, 77). We thus hypothesize that human macrophages responding to pulmonary infection may transport the virus across organs. However, the murine macrophages expressing hACE2 were clearly less capable of disseminating the virus, raising the possibility that other human cell types in our HIS mice are needed for this function. Our model is uniquely positioned to address this previously-inaccessible aspect of SARS-CoV-2 infection.

A subset of the hACE2 KI NSG mice used in our experiments also expressed an HLA-A2 transgene. HLA-A2 enables presentation of viral peptides to human CD8+ T cells (78) and its transgenic expression in mice could facilitate human T cell-murine APC and target cell interactions following infection. However, we did not detect significant differences in viral RNA levels between mice with and without the HLA-A2 transgene, and HLA expression alone may be insufficient to overcome incomplete molecular interactions between species in pathways mediating adhesion, co-stimulation and cytokine responses. Nevertheless, in the snRNA-seq analysis, which was performed on hACE2 KI HLA-A2 Tg mice, interactions between human immune cells and murine lung cells resulted in a broadening of DEGs across the murine lung cell compartment in infected HIS mice compared to immunodeficient mice, demonstrating cross-talk between the murine lung cells and human immune cells.

While observations in other SARS-CoV-2 mouse models have demonstrated lung pathology (21, 36–41), the model described here is the first to empower long-term study in animals with human immune systems that includes human fetal thymus tissue, which is essential for normal human T cell development and selection, with robust naïve T cell reconstitution in the periphery (27, 29, 33). In previous HIS mouse studies of SARS-CoV-2 infection, the human T cells had developed in the murine thymic rudiment (28, 79), which only allows slow peripheral reconstitution of human T cells. Furthermore, these are aberrantly selected in the murine thymus, which lacks normal structure, resulting in prominent autoimmune responses (27, 29) complicating interpretation of responses to infection. Lung pathology in our model included the presence of microthrombi, fibrin formation, and thickening of alveolar septae, resembling acute and long-term lung injury observed in COVID-19 patients (41, 80, 81). Fibrosis scoring increased between 2 and 8 weeks post-infection, recapitulating the sustained pulmonary fibrotic response observed in PASC. Single nucleus RNA-sequencing revealed a coordinated, tissue-wide response to infection involving chemokine production by murine alveolar macrophages, replicating the dysregulated immune signature observed in COVID-19 patients (59, 82). Together, these data indicate a coordinated immune response between human and mouse cells that drives long-term pathology.

Comparison of acutely infected and uninfected HIS mouse lung samples confirmed the enrichment of anti-viral immune response pathways, including NF-kB signaling, TLR signaling, and inflammasome activation pathways in human cells in infected mice.

These pathways have been shown to initiate viral clearance through the production of IFN I and III and pro-inflammatory cytokines, which further activate ISGs to stop viral replication while activating dendritic cells and natural killer cells that prime the adaptive immune response(83–86).

Multiple clinical studies have documented ongoing T cell responses in PASC patients months after initial infection, including SARS-CoV-2-specific CD4+ T cells with exhaustion phenotypes and persistent interferon-stimulated gene expression (87–92). A prominent T cell response was detected in Hu/Hu mice at early and late timepoints.

These included SARS-CoV-2 peptide-specific T cell responses in acute and long-term infected mice and the presence of SARS-CoV-2-specific TCR sequences. Long-lived S protein-specific CD4+ T cells with cytotoxic transcriptomic profiles driven by GZMA have been detected at 3-4 years post-infection in humans (93).Consistently, infected Hu/Hu mice exhibited a distinct shift towards mature CD4+ TEMRA cells with a unique cytotoxic transcriptional program and reduction in naïve CD4+ T cells that echoed this profile. At week 8 post-infection, both CD4+ and CD8+ T cells demonstrated upregulation of IFN response and JAK-STAT signaling, indicative of persistent immune activation. The temporal dynamics observed in our model bear striking resemblance to features of PASC, including persistent viral RNA at 8 weeks post-infection, ongoing T cell activation including cytotoxic CD4+ cells, and a delayed IFN response, with prominent IFN and JAK/STAT activation at week 8 in animals with human T cells.

However, the innate response characterized by upregulation of interferon-inducible and inflammasome-related genes at early timepoints was markedly impaired by the presence of T cells. These findings are consistent with studies showing that adaptive immunity can temper innate responses to viral infection through decreased levels of inflammatory cytokines (94–99). Given the failure of this T cell response to completely eliminate detectable viral RNA or protein even by 8 weeks and in view of the importance of Type I and III interferons in controlling the severity of COVID-19 (100–102), these results suggest a model wherein an ineffective T cell response fails to clear virus, which propagates virus-induced inflammation environment in both the short and the long term, providing a possible mechanism for PASC. Consistent with this interpretation, clearance of the virus in brain tissue by 8 weeks was associated with an absence of detectable inflammatory gene upregulation. The resulting overall similar outcomes in Hu/Hu and Hu mice are consistent with human studies that have failed to show a clear protective effect of T cells against severe infection in unvaccinated individuals. Indeed COVID-19 has not been consistently shown to be more severe in patients who receive immunosuppressive medications following transplantation or for treatment of cancer or autoimmune disease (103–105).

Our studies revealed a prominent antiviral immune response in the brain and heart of both acutely infected and long-term infected hACE2 KI NSG HIS mice. IFN-γ and Type I IFN responses were enriched in both infected brain and heart tissue at acute timepoints compared to uninfected controls, with strong upregulation of TBK1 and OAS3 in hearts of infected mice. OAS gene products sense dsRNA viruses and play a critical role in establishing innate immunity through IFN responses. OAS genes have been found to be highly expressed in infected cardiomyocytes and linked to COVID-associated heart failure (63). Loss-of-function variants of OAS have been associated with severe acute COVID-19 (66). TBK1 is also involved in Type I IFN signaling and inhibition of TBK1 attenuated lung inflammation in a SARS-CoV-2 mouse model (62). Enriched genes in the heart during acute infection also included CXCL8, IL5, IL6, and IL15, which have been implicated in COVID-19 cytokine storm (44). OAS1 was also upregulated in acutely infected brain samples, indicating sensing of viral RNA and an associated IFN response. Thus, our model faithfully replicates several aspects of the multi-organ pathology of SARS-CoV-2.

Several limitations of the hACE2 KI HIS mouse model should be considered when interpreting these findings. First, many human cytokines (e.g. IL-3, IL5, GM-CSF, M-CSF, interferons) do not cross-react well with murine receptors and vice versa, potentially limiting the spectrum of human immune effects in a murine microenvironment (22). Additionally, incompatible adhesion interactions and MHC disparities between the selecting human thymus and murine cells in the periphery present further limitations in mouse-human cellular communication. Our snRNA-seq data suggested communication between human immune and mouse lung cells in hACE2 KI HLA-Tg HIS mice, but we did not compare these mice to a group of hACE2 KI HIS mice lacking HLA-A2. NSG mice also do not develop normal peripheral lymphoid structures due to the absence of functioning lymphoid tissue inducer cells, affecting normal cell trafficking. Another limitation is that our studies were confined to a single, early SARS-CoV2 strain, and altered characteristics of later mutated strains might result in significantly different outcomes in our model. Additionally, while viral RNA and protein persisted in multiple tissues up to 8 weeks post-infection in the hACE2 KI HIS mouse model with the WA1 strain of SARS-CoV-2, we did not directly assess replication-competent virus using plaque assays or subgenomic RNA analysis at late time points. Therefore, these findings should be interpreted as evidence of prolonged viral persistence signatures rather than definitive evidence of ongoing productive infection. This model is uniquely positioned to assess differences in short- and long-term pathologies induced by different variants of the virus.

In summary, our novel hACE2 KI NSG HIS mouse provides new opportunities to explore COVID-19 and PASC pathogenesis and treatments. By combining physiologic hACE2 expression with human immune systems including T cells that undergo normal selection in a human thymus, our model avoids many of the limitations of other mouse models for the study of COVID-19. We have observed persistent viral RNA and protein across multiple organs for up to 8 weeks along with inflammation and features of PASC. We have demonstrated that human immune cells promote the dissemination of virus to extrapulmonary organs and increase the level of pulmonary infection. Persistent viral RNA was associated with strong, long-lasting anti-viral immune responses in the heart while clearance of virus over time resolved inflammation in the brain. Our model uniquely recapitulates both prolonged viral persistence signatures and a human adaptive immune response in the context of physiologic hACE2 expression, providing new opportunities to explore PASC pathogenesis and evaluate therapeutic interventions targeting the virus-immune interface.

## Materials and Methods

### hACE2 KI and hACE2 KI HLA-A2 transgenic mouse generation

hACE2 KI NSG mice were generated under contract with The Jackson Laboratory on a NOD.Cg-Prkdc^scid^ Il2rg^tm1Wjl^/SzJ (Strain# 005557) background. The human ACE2 cDNA was inserted at the start codon of mouse ACE2 in exon 2 via CRISPR/Cas9 by using guide RNAs for exon 2 (Fig. S9). Embryo microinjection was performed along with donor DNA plasmids to execute the knock-in. A subset of hACE2 KI HLA-A2 transgenic mice (n=17) were generated for use in this study by breeding homozygous hACE2 KI NSG mice with HLA-A2 NSG mice, obtained from The Jackson Laboratory (Strain #009617). After establishment of the lines, colonies of hACE2 KI NSG and hACE2 KI HLA-A2 Tg mice were then maintained at Columbia University Medical Center (CUMC) under specific pathogen-free conditions in our Helicobacter and Pasteurella pneumotropica-free animal facility. All animal experiments were performed with approval from the CUMC Institutional Care and Use Committee. SARS-CoV-2-infected mice were housed in an ABSL-3 facility with approval from the CUMC Institutional Animal Care and Use Committee and CUMC Environmental Health and Safety.

### Validation of hACE2 expression

DNA and RNA were isolated from hACE2 KI NSG mice using the Qiagen DNA/RNA Mini Kit (Cat# 80204). The presence of the hACE2 insert was validated in genomic DNA using hACE2 F2 primer (TGGTCCAGCAGCTTGTTTACT) and hACE2 R2 primer (GGCAGTCACTCATCCTCACA). The resulting PCR product was run on a 1.2% agarose gel and demonstrates the expected insert size for hACE2 of 3081bp.

A 2-step qPCR for RNA-level hACE2 expression was performed using RNA isolated from lung, brain, brain stem, heart, liver, spleen, reproductive organs, kidney, stomach, duodenum, jejunum, ileum, and colon tissue harvested from hACE2 KI NSG mice.

Reverse transcription was performed using Superscript III Reverse Transcriptase (Thermofisher Cat# 18080093) followed by cDNA synthesis using hACE2 F primer (CGAAGCCGAAGACCTGTTCTA) and hACE2 R primer (GGGCAAGTGTGGACTGTTCC). Fold change of hACE2 expression in hACE2 KI NSG mouse tissue was calculated against hACE2 expression in human lung tissue, using hACE2 expression in A2-transgenic NSG liver tissue as a negative control alongside no-template and no-RT controls.

IHC staining was performed by the Molecular Pathology Shared Resource facility at Columbia University Medical Center. Lung and intestinal tissues from hACE2 KI NSG mice were fixed in 4% paraformaldehyde for 24 hours, embedded in paraffin, and sectioned into 5 um blank slides. Slides were stained with hematoxylin and eosin for morphology check. IHC staining was performed on 5 μm-thick FFPE sections with rabbit anti-hACE2 antibodies (Abcam ab108209) to detect hACE2 expression. Antigen retrieval was performed in 10 mmol/L citrate buffers (pH 6.0) with a pressure cooker.

Quenching of the endogenous peroxidase activity was done with 3% hydrogen peroxide in PBS-T for 10 minutes at room temperature. Slides were incubated with primary antibody anti-hACE2 1:1000 for 2 hours at room temperature, followed by incubating with anti-rabbit polymer (DAKO #K4003) for 45 minutes at room temperature. Slides were visualized with HRP catalyzed chromogen “diaminobenzidine”, then counterstained with hematoxylin, dehydrated, cleared in xylene, and mounted with Permount. Imaging was done with Leica SCN400.

### Generation of human immune systems

All human fetal tissues (gestational age 17-21 weeks) were obtained from Advanced Bioscience Resources. Fetal thymus and CD34+ hematopoietic stem cells were prepared as previously described (32). Briefly, fetal thymi were processed into 2x2x2mm fragments and cryopreserved using 10% DMSO and 90% human AB serum (Gemini Bio-Products). To isolate CD34+ hematopoietic stem cells, fetal liver cell suspensions were generated by density gradient separation using Histopaque 1077 (Sigma Aldrich). Then, CD34+ cells were isolated using positive selection by magnetic-activated cell sorting with anti-human CD34+ microbeads (Miltenyi Biotec).

At 6-8 weeks of age, mice were thymectomized to remove the native mouse thymus (29). Two weeks after this procedure, the mice underwent either sublethal total body irradiation (1Gy) by an X-Ray irradiator (RS-2000, Rad Source Technologies, Inc., Suwanee, GA) or were intraperitoneally injected with busulfan (0.5mg/mL). Half of the mice were engrafted with a cryopreserved and thawed 2mm^3^ fragment of human fetal thymus under the kidney capsule. These are referred to as Hu/Hu mice. Both Hu mice, which did not receive fetal thymus, and Hu/Hu mice were then intravenously injected with 2x10^5^ magnetically isolated CD34+ cells from human fetal liver, from the same donor as the human fetal thymus.

Human cell reconstitution was monitored via bi-weekly flow cytometric analysis of mouse peripheral blood mononuclear cells (PBMCs) using an LSRII (BD) or Cytek Aurora (Cytek Biosciences) analyzer beginning at week 5 post-transplantation as previously described (106). Approximately 100ul of whole blood was collected via tail vein puncture. PBMCs were then isolated by density gradient separation using Histopaque 1077 (Sigma Aldrich) for flow cytometry. Antibody information is available in Table S8.

### SARS-CoV-2 infection

SARS-CoV-2 Virus Isolate USA-WA1/2020 was provided by the Aaron Diamond AIDS Research Center BSL-3 Core Facility at CUMC. Mice were intranasally infected with 5,000 pfu of virus (in acute infection studies) or 10,000pfu of virus (in long-term studies) in an ABSL-3 facility by trained ICM staff.

### Tissue processing

Mice were euthanized at days 7-, 14-, or week 8 post-infection, which was 14-22 weeks post-transplantation of human fetal tissue. At harvest, blood was obtained via cardiac puncture and centrifuged at 4500prm for 10 minutes in order to collect plasma. Virus was subsequently inactivated by adding 10% volume of TritonX-100 to the isolated plasma and shaking for 2 hours. Spleen, lung, brain, and heart were harvested from each mouse for further processing. Each tissue was separated for histology and RNA isolation in Trizol. A portion of the spleen was crushed to obtain a single cell suspension, which was then filtered through a 70um nylon filter. Spleens were subsequently treated with Ammonium-Chloride-Potassium (ACK) erythrocyte lysis buffer (Life Technologies). A portion of lung (<50mg) was snap frozen on dry ice and transferred to liquid nitrogen for downstream single nucleus RNA-sequencing. Remaining lung tissue was then dissociated by chopping into fine pieces (approximately 5mm^3^), resuspending in reconstituted liberase (400ug/mL), and incubated at 37C for 30 minutes before being washed. Cells from the PBMC, spleen, and lung were then counted and stained for flow cytometry. Antibody information is available in Table S8. Antibody surface staining was performed at room temperature for 20 minutes and cells were then fixed using 4% paraformaldehyde. Samples were collected on the Cytek Aurora (Cytek Biosciences) and analyzed using FlowJo software.

### Detection of virus

Using RNA extracted from lung, brain, and heart collected in Trizol at each timepoint, SARS-CoV-2 spike protein was quantified using Applied Biosystems TaqPath One Step RT qPCR Master Mix (Thermo Scientific #A15299) and TaqMan Microbe Detection Assay SARS-CoV-2 S Gene probes (Thermo Scientific #A50137 ID Vi07918636_s1) with fold change calculated against mouse GAPDH (Thermo Scientific #4331182 ID Mm99999915_g1). Nucleocapsid protein was quantified using the 2019-nCoV RUO Kit (IDT #10006713) and 2019-nCoV Positive Control (IDT #10006625).

### Histology preparation and immunohistochemistry

IHC staining was performed by the Molecular Pathology Shared Resource facility at CUMC. Lung tissues were fixed in 4% paraformaldehyde for 24 hours, embedded in paraffin, and 5 um blank slides were sectioned. Slides were stained with hematoxylin and eosin (H&E) for morphologic examination. IHC staining was performed on 5 μm-thick FFPE sections. Antibody information is available in Table S9. Antigen retrieval was performed in 10 mmol/L citrate buffers (pH 6.0) with a pressure cooker. Quenching of the endogenous peroxidase activity was done with 3% hydrogen peroxide in PBS for 10 minutes at room temperature. Slides were incubated with primary antibody for 90 minutes to 2 hours at room temperature, followed by incubation with biotinylated goat anti-rabbit IgG (H+L, Vector Laboratories, Inc. #BA_1000) at 1:200 dilution and tertiary streptavidin-peroxidase conjugate (ABC complex, Vector Laboratories, Inc. # PK-6100) or incubation for 45 minutes with anti-rabbit polymer (DAKO #K4003). Slides were visualized with HRP catalyzed chromogen “diaminobenzidine”, then counterstained with hematoxylin, dehydrated, cleared in xylene, and mounted with Permount. Slides were imaged with Leica SCN400 and IHC staining was quantified using HALO image analysis software.

### Tissue modeling for fibrosis quantification

Sixty-seven H&E-stained lung sections were digitized using an Aperio slide scanner at a resolution of 0.252 um/pixel (40x equivalent) and stored as SVS whole-slide images.

Each slide was divided into non-overlapping 224 x 224-pixel patches, retaining only patches in which tissue occupied at least 50% of the area (307,118 patches total). Patch-level feature embeddings (1,536 dimensions) were extracted using the UNI2-h vision transformer (ViT-H/14) foundation model, which was pre-trained on over 100 million histopathology images(107). To correct for batch effects introduced by tissue perfusion status, ComBat harmonization(108) was applied to the slide-level mean feature vectors using the *sva* package(109) in R (v4.5.1), with treatment group specified as a biological covariate to preserve. The corrected features were reduced to 100 principal components and clustered by MiniBatchKMeans (K = 8, batch size = 10,000, 10 random initializations, seed = 42). One cluster (C7) was identified as fibrotic tissue on the basis of differential enrichment in infected slides and visual inspection of representative patches.

For cell-level analysis, nuclei were segmented and classified by two complementary approaches. First, HoVer-Net(110), pretrained on the PanNuke dataset, was applied via TIAToolbox to classify each nucleus into one of five types: neoplastic epithelial, inflammatory, connective/fibroblast, dead/apoptotic, and non-neoplastic epithelial (816,320 cells total across 67 slides). Second, for morphometry, nuclei were independently segmented using Cellpose with default parameters (estimated nuclear diameter = 30 pixels, flow threshold = 0.4)(111). Three morphometric features—area, eccentricity, and solidity—were extracted from each segmented nucleus using scikit-image regionprops. After log-transformation of area and z-score normalization, cells were clustered by MiniBatchKMeans (K = 4) to identify morphological phenotypes, including an elongated, high-eccentricity population consistent with fibroblasts. The HoVer-Net connective/fibroblast proportion and the morphometric fibroblast cluster proportion showed concordant patterns across slides. A composite fibrosis score was computed for each slide as the mean of three z-scored components: (i) the proportion of patches assigned to the fibrotic tissue cluster (C7), (ii) the mean nuclear eccentricity, and (iii) the proportion of cells assigned to the fibroblast morphological cluster. Spatial autocorrelation of fibrotic patches was assessed by Moran’s I, and focal fibrotic hotspots were identified by the Getis-Ord Gi* statistic, both computed on a k-nearest-neighbor graph (k = 8) of patch coordinates. Fibrosis composite scores were compared between infected and control mice across five pre-specified cohort contrasts (Combined, Acute, Long-term 1, Long-term 2, Combined Long-term) using the Mann-Whitney U test. Three slides of unknown treatment status were excluded from statistical comparisons. The complete fibrosis quantification pipeline is available at https://github.com/princello/pathology-fibrosis-pipeline.

### ELISpot

Human interferon gamma (IFN-γ) ELISpot assay was performed as per the manufacturer’s instructions (Mabtech, Nacka Strand, Sweden). Briefly, after collecting splenocytes or lung mononuclear cells from SARS-CoV-2-infected Hu/Hu mice, the cells were resuspended in complete medium (RPMI-1640 with 10% heat-inactivated fetal calf serum), 1 mM sodium pyruvate, 100 U/mL penicillin, 100 μg/mL streptomycin, 10 mM HEPES (Sigma), and 10 IU/mL of recombinant human IL-2 (PeproTech, Cranbury, NJ), and rested on ice for one hour. Then, the cells were seeded at 2 x 10^5^ or 4 x 10^5^ cells/well in MultiScreen HTS Filter Plates precoated with anti-human IFN-γ antibody (15 μg/mL; clone 1-D1K; Mabtech). Test wells were then supplemented with overlapping 17-mer peptides spanning S-protein, N-protein, M-protein, and E-protein (each at 1 μg/mL; BEI Resources, Manassas, VA). Negative control wells lacked peptides. Positive control wells contained Staphylococcal enterotoxin B (1 µg/mL; Sigma). The cell cultures were incubated for 24 hours at 37°C in a 5% CO_2_ incubator. Plates were then washed five times with PBS (Sigma-Aldrich) and incubated for 2 hours at room temperature with HRP-conjugated anti-human IFN-γ antibody (1 μg/mL; clone 7-B6-1; Mabtech). Plates were then washed a further five times with PBS and developed for 10 minutes with BCIP/NBT Substrate (Mabtech). The color development (reaction) was stopped by washing extensively in deionized water, and the plates were allowed to dry. Finally, the spots were counted using an ELISpot reader. All assays were performed in duplicate.

### Nanostring gene expression analysis

Nanostring was performed by the Human Immune Monitoring Core at CUMC and the Albert Einstein College of Medicine Immune Monitoring Core using an input of 100ng of RNA. Samples were run using the nCounter Host Response panel with the nCounter Coronavirus Panel Plus Spike-in included for acute lung samples only. Raw data was exported into R for further analysis. Gene expression was normalized using the *NACHO* R package (V2.0.6) (112). *DeSeq2* (V1.46.0) was used to detect differentially expressed genes among experimental conditions (113). Genes were detected as differentially expressed using an adjusted *p*-value cutoff of <0.05.

Gene set enrichment analysis (GSEA) was performed using the *fgsea* R package (V1.32.4) (114). The human hallmark pathways and canonical pathways gene sets were used, obtained from the Molecular Signatures Database (MsigDB) (60, 61). Statistical analysis from *DeSeq2* was used as the input. Graphical representations for differentially expressed genes and GSEA analysis were generated using R packages *EnhancedVolcano* (V1.24.0) and *ggplot2* (V3.5.1) (115, 116).

Cross-reactivity of the Nanostring nCounter Host Response panel (773 human-targeted probes) with murine transcripts was evaluated using an NSG control lung sample (animal 697) that lacked human immune cells, such that any detectable signal in this sample reflects cross-hybridization of human probes to murine transcripts. Raw counts were first normalized using the internal positive spike-in controls (geometric mean scaling) via *ncountr* v0.2.0. For each probe, a cross-reactivity ratio was computed as the positive-control-normalized NSG-697 count divided by the median count across four Hu/Hu control samples, which represent genuine human signal. Probes were classified as reliable (ratio < 0.10, n = 436, 56%), moderate (0.10 ≤ ratio < 0.50, n = 200, 26%), or unreliable (ratio ≥ 0.50, n = 137, 18%). For the 38 IFN/JAK-STAT pathway genes represented in the panel, 34 (89%) were classified as reliable. To present probe reliability in a more interpretable form, we additionally report the human signal fraction for each gene, defined as ‘max(0, 1 − cross-reactivity ratio)’; this quantity represents the proportion of the Hu/Hu signal attributable to genuine human transcript and ranges from 0 (mouse-dominated) to 1 (purely human). Three IFN genes (IRF1, IFNB1, DDX58) had cross-reactivity ratios ≥ 0.50; these were flagged in supplementary materials but retained in pathway-level analyses, as their exclusion did not alter the overall IFN pathway enrichment conclusions. Full per-probe cross-reactivity ratios, reliability classifications, and human signal fractions for all IFN genes are provided in Supplementary Table S10.

### Single nucleus RNA-seq

Single nucleus RNA-sequencing was performed on a subset of mice at 8 weeks post-infection. Tissue was prepared as described by Melms et al. with minor modifications(117). Snap-frozen tissues were allowed to equilibrate on wet ice. Tissues were resuspended in Tween with salts and Tris buffer (TST) and mechanically dissociated using fine scissors. Tissues were then pipetted up and down for further dissociation. Samples were then filtered through a 40um filter and washed with ST buffer. Nuclei were then counted and 14,000-16,000 were then loaded into each channel on a Chromium controller using Chromium Next GEM Single Cell 5 v2 reagents (10X Genomics, Pleasanton, CA). Single-nucleus RNA-seq libraries were prepared per the manufacturer’s instructions. Libraries were analyzed and quantified using TapeStation D1000 screening tapes (Agilent, Santa Clara, CA) and Qubit HS DNA quantification kit (Thermo Fisher Scientific). Libraries were pooled and sequenced on a NovaSeq 6000 at the Columbia Genome Center.

### T cell receptor sequencing

Samples were prepared using the Chromium Single Cell Human TCR Amplification Kit (10X Genomic, Pleasanton, CA) per the manufacturer’s instructions, using the same sample input as in the single nucleus RNA-sequencing. Libraries were run alongside the single nucleus RNA-seq samples on the NovaSeq 6000 at the Columbia Genome Center.

### Analysis of sequencing data

#### snRNA-seq preprocessing and integration

Sequencing reads were processed with the 10x Genomics Cell Ranger pipeline. In samples containing both human and mouse cells, nuclei were assigned to a species when more than 80% of uniquely mapped reads aligned to the respective genome; nuclei not meeting this threshold were excluded. Quality control filtering removed nuclei with fewer than 200 detected genes or with mitochondrial read fractions exceeding 10%. Gene expression counts were normalized to 10,000 counts per cell and log-transformed (log1p). The top 4,000 highly variable genes were selected using the *Seurat v3* method with batch-aware estimation (batch_key = sample).

Samples were integrated using *scVI* (118) with a negative binomial likelihood, 256 hidden units, 4 layers, a 24-dimensional latent space (dropout rate = 0.2), and gene-batch dispersion, conditioning on sample, condition, and mouse type. Cell type annotation was performed with *CellTypist*(*119*): mouse nuclei were classified against the LungMAP MouseLung CellRef v1.1 reference(120), yielding 16 major cell types across 49,772 cells from 16 samples; human nuclei were classified against the Immune_All_High reference, yielding 8 major cell types across 4,401 cells from 10 samples (6 Hu/Hu, 4 Hu).

#### Differential expression

Differential gene expression between infected and uninfected groups was assessed at two levels. For pseudobulk analysis, raw counts were aggregated per sample per cell type, and differentially expressed genes were identified using *pyDESeq2* with default parameters(121). For single-cell-level analysis, the Wilcoxon rank-sum test was applied via scanpy.tl.rank_genes_groups(122).

#### CD4⁺ T cell differential expression and exclusion of sex-chromosome confounders

Human CD4⁺ T cells (n = 271 from six Hu/Hu mouse samples) were identified from the integrated human snRNA-seq dataset by selecting cells with *CD4* expression and no detectable *CD8A* or *CD8B* expression. Differential gene expression between infected (n=3) and uninfected (n=3) samples was computed at the single-cell level using the Wilcoxon rank-sum test implemented in ‘scanpy.tl.rank\_genes\_groups’, with Benjamini-Hochberg multiple-testing correction across all 14,699 genes. Initial inspection of the volcano plot revealed that the most statistically significant genes in both directions were driven by donor sex imbalance: Y-chromosome genes (UTY, USP9Y, RPS4Y1, KDM5D, DDX3Y, EIF1AY) dominated the down-regulated hits, and the X-inactive specific transcript XIST together with the X-escape gene KDM6A dominated the up-regulated hits. To obtain a biologically interpretable view of the infection response, we annotated all 14,699 tested genes with their chromosome of origin using a Biomart query against the Ensembl GRCh38 assembly (R biomaRt/gseapy.Biomart) and filtered out all 439 genes annotated to the X or Y chromosome.

#### Pathway and transcription factor analysis

Pathway activity was inferred using *decoupleR*(*123*). Transcription factor activity was estimated by the univariate linear model (ULM) method with the CollecTRI regulon database. Pathway-level scores were obtained from the PROGENy model and the MSigDB Hallmark gene set collection.

#### CD4 T cell subset analysis

Human CD4+ T cells (271 cells) were scored for five functional subsets using scanpy.tl.score_genes with the following marker gene sets: TEMRA (CX3CR1, GZMB, PRF1, GZMA, KLRG1, NKG7, GNLY, FGFBP2), effector memory (CCL5, GZMK, OMES, IL7R), central memory (CCR7, SELL, LEF1, TCF7, IL7R), naive (CCR7, SELL, LEF1, TCF7, CD27), and regulatory T cells (FOXP3, IL2RA, CTLA4, TIGIT, IKZF2). Pseudotime ordering of T cells was computed with Palantir(124) using diffusion maps on the *scVI* latent embedding.

#### CD8⁺ T cell analysis

Human CD8⁺ T cells were identified from the Hu/Hu T cell population (1,628 cells) by selecting cells with CD8A or CD8B expression > 0 and no detectable CD4 expression, yielding 131 cells (79 infected, 52 uninfected) across 6 Hu/Hu samples. CD8-specific functional subset signatures were scored using ‘scanpy.tl.score\_genes’ with the following curated gene sets: Naive (CCR7, SELL, LEF1, TCF7, CD27, IL7R), central memory (CCR7, SELL, TCF7, IL7R), effector memory (GZMK, CCL5, EOMES, IL7R), TEMRA (CX3CR1, GZMB, PRF1, GZMA, KLRG1, NKG7, GNLY), and tissue-resident memory/TRM (CD69, ITGA1, CXCR6, RUNX3, ITGAE). Scores were compared between infected and uninfected conditions using the Mann-Whitney U test at both single-cell and pseudobulk (per-sample mean) levels.

Pathway activity was inferred using decoupler: Hallmark enrichment by univariate linear model (ULM) and PROGENy pathway activity by multivariate linear model (MLM), applied to the Wilcoxon z-score vector (single-cell analysis) or pyDESeq2 Wald statistic (pseudobulk analysis).

Statistical power limitations: Only 131 CD8⁺ T cells (6–49 per sample) were detected, limiting single-cell differential expression analysis. Despite this per-gene limitation, pathway-level analyses (which aggregate signal across hundreds of genes) detected significant IFN/JAK-STAT activation (Hallmark IFN-alpha, IFN-gamma; PROGENy JAK-STAT), and gene-set scoring detected significant CD8⁺ subset changes (Naive and CM down, TEMRA and TRM up) in infected mice.

#### Cell composition analysis

Changes in cell type proportions were assessed by *scCODA*(*125*) using Markov chain Monte Carlo inference, with cell types considered credibly altered at an inclusion probability above 0.9.

#### Cell-cell communication

Ligand-receptor interactions were inferred with *LIANA*(*126*) using the rank_aggregate method and the consensus (human) or mouse consensus (mouse) interaction resource.

#### TCR analysis

SARS-CoV-2-specific TCR detection: T-cell receptor CDR3 sequences assembled by TRUST4 v1.1.8 from 10x VDJ data were matched against a combined SARS-CoV-2 reference constructed by pooling human CDR3β and CDR3α sequences from VDJdb (antigen.species = "SARS-CoV-2", HomoSapiens), McPAS-TCR (Pathology = "COVID-19"), and IEDB (SARS-CoV-2 epitope-specific T cell receptors), yielding 21,653 unique CDR3β and 4,188 unique CDR3α reference sequences (127–129). Assembled contigs were indexed with *scirpy*(130). For each sample contig we computed three independent match flags: (i) *exact*, the CDR3 amino-acid sequence appeared verbatim in the SARS reference set; (ii) *Hamming* ≤ *1*, the CDR3 was the same length as a reference sequence and differed at exactly one position; (iii) *GLIPH2 motif*, the sample CDR3 shared a short amino-acid motif with the SARS reference set that had been previously identified as enriched in SARS-specific relative to non-SARS-specific human TCRs.

GLIPH2 motif enrichment was performed separately for each chain following a simplified version of the Huang et al. (2020, *Nature Biotechnology*) algorithm (131). Conserved N- and C-terminal anchor residues were trimmed from each reference CDR3 (three amino acids from each end), and all 4-mers were extracted from the trimmed cores. Each 4-mer was tested for enrichment in the SARS reference set compared to a background of all non-SARS human CDR3 sequences from VDJdb (78,422 CDR3β, 34,444 CDR3α) using Fisher’s exact test on counts of unique sequences containing the motif. For CDR3β we used strict criteria — minimum 5 SARS sequences containing the motif, fold enrichment ≥ 10× compared to background, and Fisher p ≤ 10⁻¹ — which yielded 10 enriched motifs (APQE, AQNT, DANT, DQNT, EQNT, HNTG, MNTG, PGPP, QNTG, SLED). For CDR3α we used relaxed criteria (minimum 3 SARS sequences, fold ≥ 3, p ≤ 10⁻³) necessitated by the smaller SARS TRA reference pool; this yielded 6 enriched motifs (FSGS, GGQK, NEED, NKED, NRED, PTGT). A sample contig was flagged as a GLIPH2 match if its core contained any enriched motif.

Enrichment of SARS-CoV-2-matching TCRs in infected relative to uninfected samples was tested with two complementary approaches: (i) Fisher’s exact test on pooled contingency tables across all contigs from each condition (the "pooled contig-level" test), which asks whether a randomly drawn contig is more likely to match a SARS reference in infected than in uninfected samples and is sensitive to clonal expansions within a single sample; and (ii) Mann-Whitney U on per-sample match rates treating each mouse as an independent experimental unit (the "per-sample" test), which asks whether infected mice show higher rates than uninfected mice and is conservative at small sample sizes (n = 3 vs 3 gives a minimum achievable two-sided p ≈ 0.1). For TRA analyses involving Hamming ≤ 1, the two tests give discordant directions (Fisher infected > uninfected, Mann-Whitney uninfected > infected), a Simpson’s paradox driven by the disproportionately large TRA library of sample 678 (572 contigs, 40.9% match rate) dominating the pooled infected-condition contigs.

### Statistics

Statistical analysis was performed using GraphPad PRISM version 10 software (San Diego, CA). Unpaired t-tests and ordinary one-way ANOVA were used to determine significance between independent groups. Data in graphs are expressed using the mean ± SEM. A p-value of <0.05 was considered statistically significant. For computational analyses, differential gene expression in snRNA-seq data was assessed using *pyDESeq2*(*121*) for pseudobulk comparisons and the Wilcoxon rank-sum test for single-cell comparisons, with Benjamini-Hochberg false discovery rate correction applied to all multiple-testing scenarios. Cell composition changes were evaluated by *scCODA*(*125*) using Hamiltonian Monte Carlo sampling. TCR enrichment was tested by Fisher’s exact test on pooled contingency tables. Fibrosis composite scores and cell type proportions were compared between groups using the Mann-Whitney U test.

Cross-modality and cross-species correlations were assessed by Spearman’s rank correlation. All computational statistics were performed in Python 3.11 using *scipy* (v1.11)(132), *statsmodels* (v0.14) (https://www.statsmodels.org/stable/index.html), and the analysis packages listed above.

### Study approval

The acquisition of human tissues and animal study were reviewed and approved by Institutional Review Board and Animal Care and Use Committees at Columbia University respectively.

## Data availability

Single-nucleus RNA-seq and TCR-seq data have been deposited in the Gene Expression Omnibus (GEO) under accession [GSEXXXXX]. Nanostring nCounter raw data (RCC files) are available upon request from the corresponding author.

Code for the computational histopathology fibrosis quantification pipeline is available at https://github.com/princello/pathology-fibrosis-pipeline. Code for Nanostring nCounter analysis is available at https://github.com/princello/ncountr. Analysis scripts for snRNA-seq, TCR, and cross-platform integration are available upon request.

## Author Contribution

AW designed and performed experiments, analyzed data and wrote the manuscript. ND, CP and MKM designed and performed experiments. VA, KL, YS, GZ, XD, HWL and MT performed experiments. CC performed ABSL3-associated mouse infection experiments. ZW performed sequencing analysis and analyzed lung fibrosis through generation of a tissue analysis model. AS performed histology analysis. MS directed the research and edited the manuscript. All authors commented on previous versions of the manuscript and approved the final manuscript.

## Funding

This research was supported by National Institute of Health grants 5T32AI106711-09 and 3R01AI138547-03S1.

## Acknowledgements

Research reported in this publication was performed in the CCTI Flow Cytometry Core, supported in part by the Office of the Director, National Institutes of Health under awards S10RR027050. The authors also acknowledge the contributions of the CCTI Human Immune Monitoring Core, the Columbia University HICCC Molecular Pathology Shared Resource, and the Albert Einstein Immune Monitoring Core. The content is solely the responsibility of the authors and does not necessarily represent the official views of the National Institutes of Health. Figures 1a and 2a were created with BioRender.com.

---

**Figure S1:**
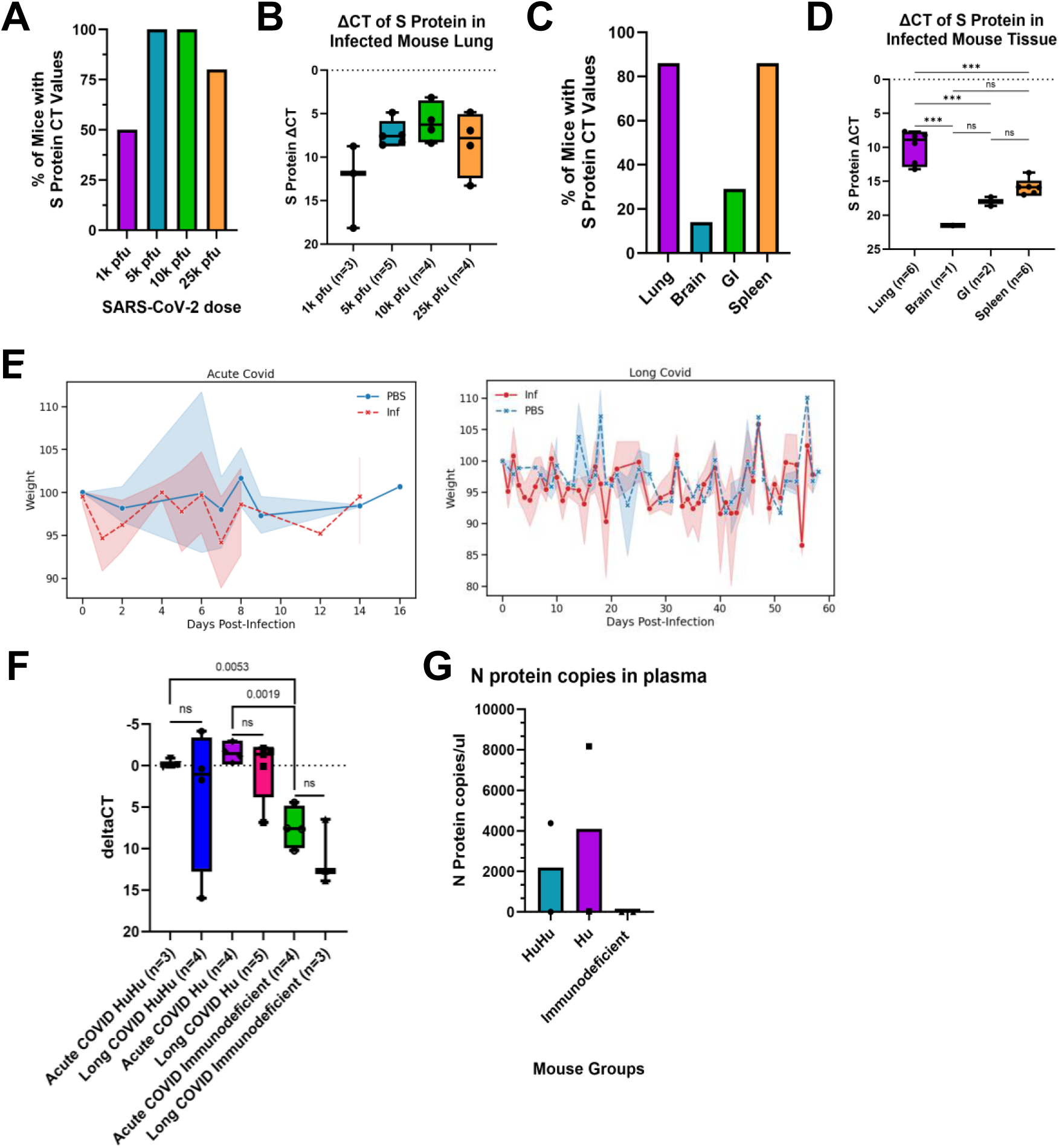
**a.** Percentage of non-humanized hACE2 KI NSG mice demonstrating SARS-CoV-2 spike protein-specific qPCR CT values intranasally infected with either 1k, 5k, 10k, or 25k pfu of WA1 at day 3 post-infection. PBS control mice (not shown) did not exhibit spike protein-specific CT values. *n*=5. **b.** Levels of viral spike protein RNA detected in the native mouse lung for all mice across all doses. ΔCT values of SARS-CoV-2 spike protein-specific qPCR were determined by normalizing the spike protein values against mouse GAPDH. There were no significant differences between groups. Box and whisker plots show min to max values. Individual values for each mouse are shown. **c.** Percentage of non-humanized hACE2 KI NSG mice intranasally infected with 5k pfu of WA1 demonstrating SARS-CoV-2 spike protein-specific qPCR CT values across multiple tissues at day 7 post-infection. PBS control mice (not shown) did not exhibit spike protein-specific CT values. *n* = 7. **d.** Levels of viral spike protein RNA detected in tissue at day 7 post-infection after infection with 5k pfu of WA1. ΔCT values of SARS-CoV-2 spike protein-specific qPCR were determined by normalizing the spike protein values against mouse GAPDH. There were no significant differences between groups. Box and whisker plots show min to max values with adjusted p-values from One Way ANOVA multiple comparisons displayed. Individual values for each mouse are shown. ***p < 0.001. **e.** Linear mixed model showing the percent weight change during the course of infection compared to baseline weight in both acute (left) and longterm (right) infection cohorts. 95% confidence intervals are shown for each group. Acute: Infected *n* = 29, Uninfected *n* = 18. Long COVID Infected *n* = 22, Uninfected *n* = 11. Infected mice in the long COVID cohort show significant weight loss compared to PBS controls. *p-*value = 0.03. There is no difference between groups in the acute infection cohort. **f.** Levels of viral spike protein RNA detected in the native mouse lung for all infected mice across both acute and long-term timepoints. ΔCT values of SARS-CoV-2 spike protein-specific qPCR were determined by normalizing the spike protein values against mouse GAPDH. Individual values for each mouse are shown with *n* denoted in group name. *p-*values between groups denoted on graph, as determined by unpaired t-tests between groups. **g.** SARS-CoV-2 nucleocapsid protein RNA copies per microliter, as determined by nucleocapsid protein-specific RT-qPCR using RNA isolated from plasma. Copy numbers determined using a standard curve from provided nucleocapsid standards. Hu/Hu *n* = 2, Hu *n* = 2, Immunodeficient *n* = 2.

**Figure S2.**
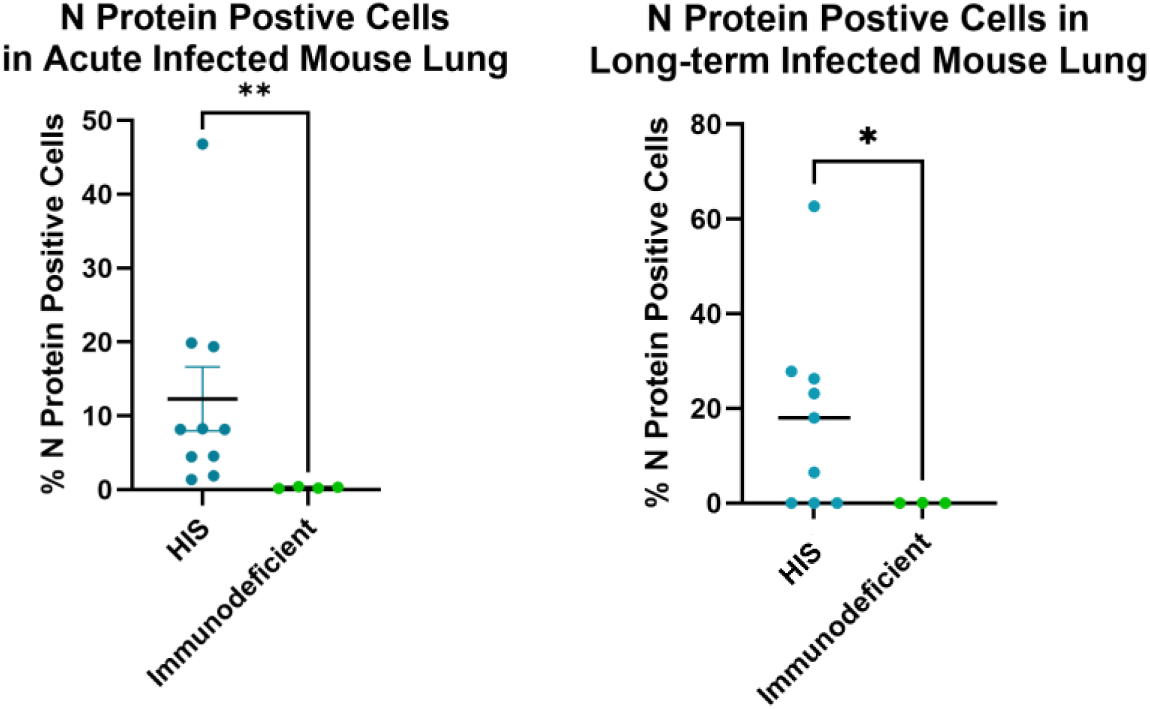
Percentage of SARS-CoV-2 nucleocapsid protein positive cells in infected HIS and immunodeficient mouse lung tissue calculated using HALO imaging software. Acute mouse (left) statistical significance was analyzed by Mann-Whitney test. *p*-value = 0.0020. HIS *n* = 10, Immunodeficient *n* = 4. Long-term mouse (right) statistical significance was analyzed by Welch’s t test. *p*-value = 0.0272. HIS *n* = 9, Immunodeficient *n* = 3.

**Figure S3.**
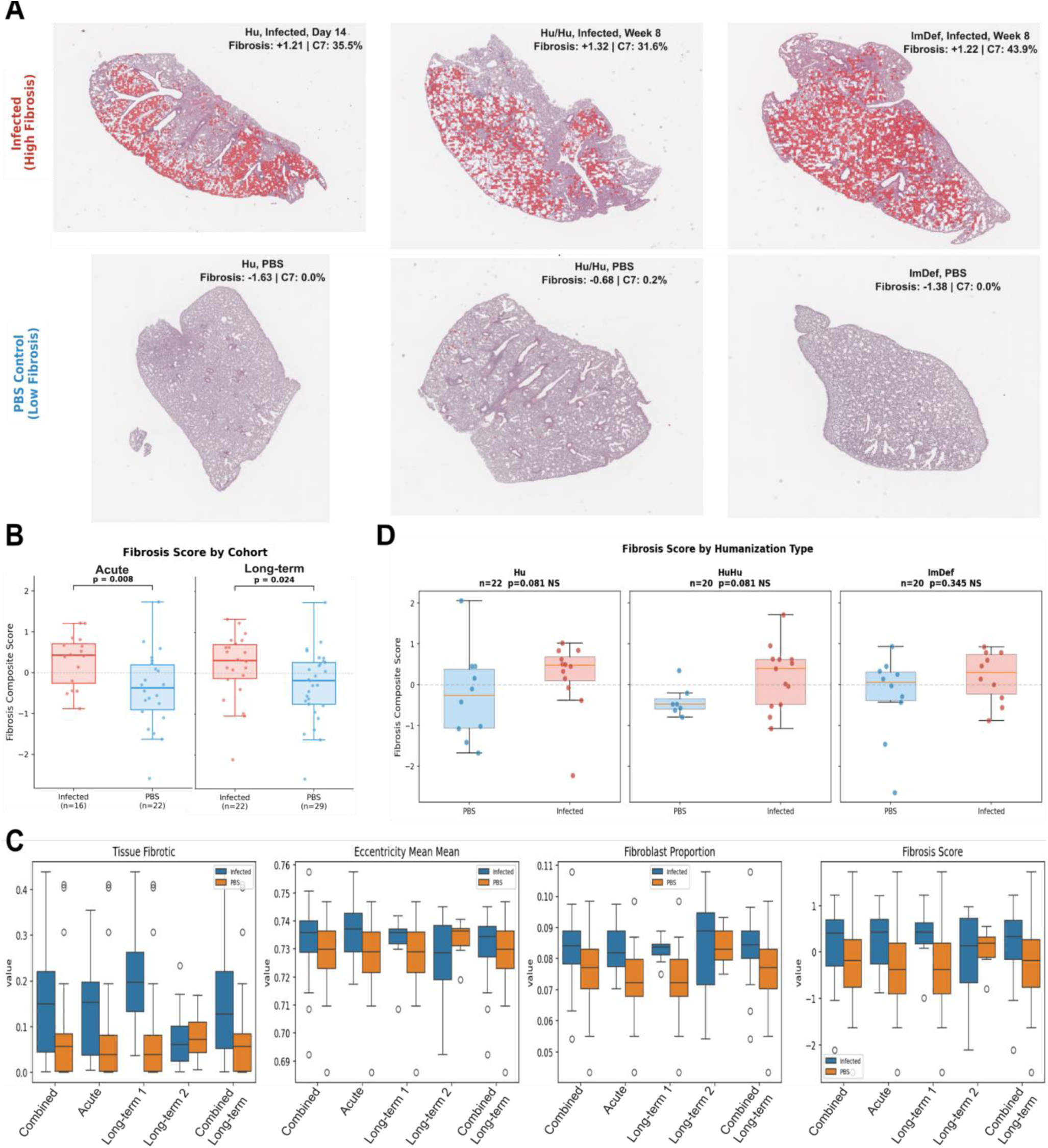
**a.** Whole-slide fibrotic cluster overlay: C7 patches (red) mapped onto representative infected (top; Hu Day14, Hu/Hu Wk8, immunodeficient Wk8) versus PBS control (bottom; Hu, Hu/Hu, immunodeficient) lung sections, demonstrating widespread fibrotic cluster distribution in infected lungs (C7 = 31.6-43.9%) compared to near-absence in controls (C7 = 0.0-0.2%). **b.** Fibrosis composite scores stratified by cohort: Acute (p = 0.008), Long-term (p = 0.024). **c.** Individual score components of the composite fibrosis score, stratified by cohort. Fibroblast proportion acute *p-value =* 0.007, tissue fibrotic proportion Long-term 1 *p-value* = 0.005 **d.** Fibrosis composite score, stratified by humanization status.

**Figure S4.**
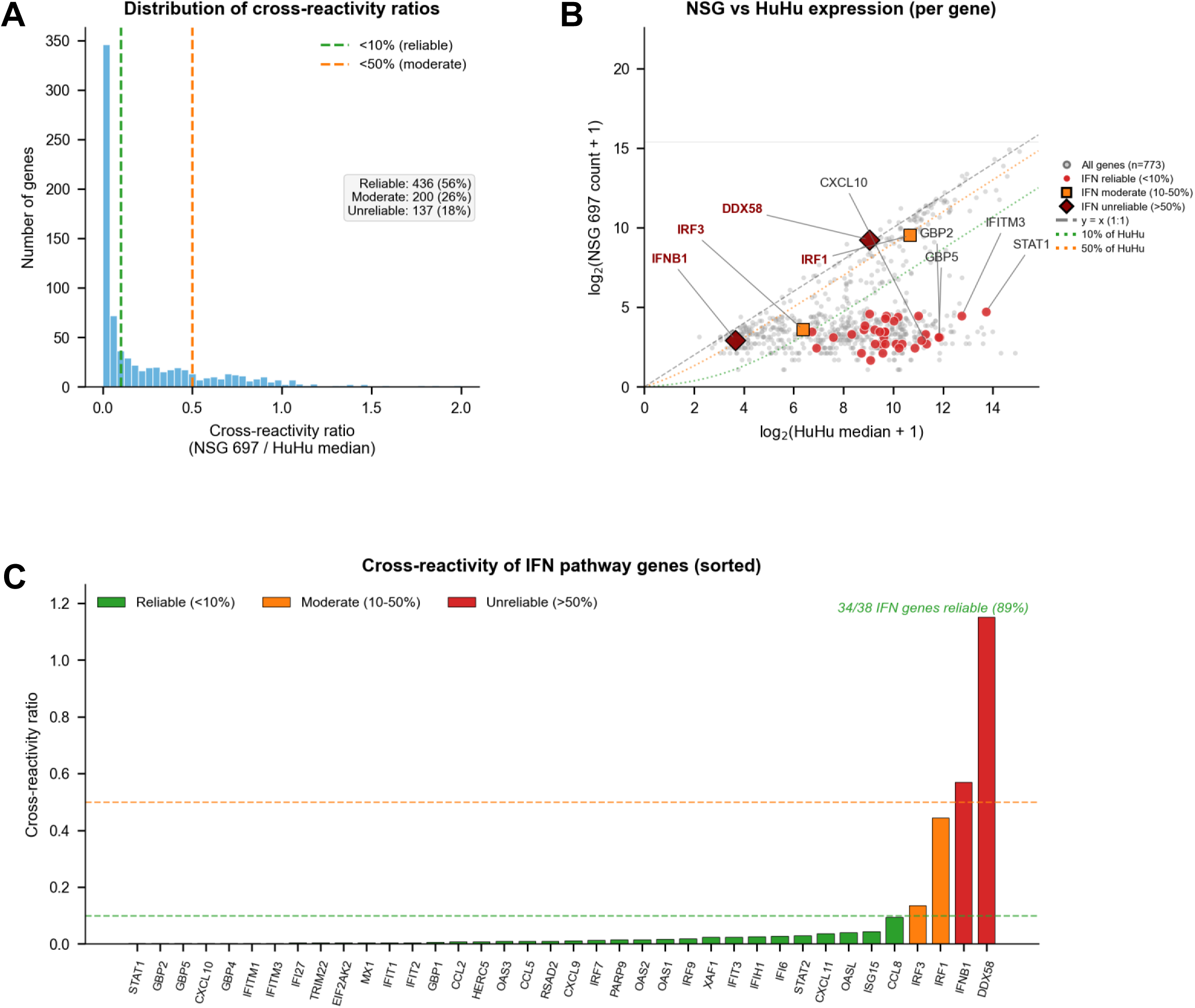
Cross-reactivity of the 773-gene human Nanostring nCounter Host Response panel was evaluated using an NSG control lung sample (animal 697), which lacks human immune cells and therefore produces signal only from murine transcripts that cross-hybridize to the human-targeted probes. For each gene, a cross-reactivity ratio was calculated as the positive-control-normalized count in the NSG sample divided by the median normalized count across four Hu/Hu samples. Probes were classified as *reliable (ratio < 0.10), moderate (0.10 ≤ ratio < 0.50), or unreliable (ratio ≥ 0.50).* **(a)** Distribution of cross-reactivity ratios across all 773 probes in the panel. Dashed lines indicate the 10% (green) and 50% (orange) thresholds; 436 genes (56%) were classified as reliable, 200 (26%) as moderate, and 137 (18%) as unreliable. **(b)** Per-gene comparison of log₂-transformed NSG-697 counts versus the log₂-transformed Hu/Hu median counts. Grey dots denote all 773 panel genes; red circles, orange squares, and dark-red diamonds highlight the 38 IFN pathway genes by reliability class. The black dashed line is the 1:1 reference (NSG = Hu/Hu); dotted green and orange lines indicate cross-reactivity ratios of 10% and 50% relative to the HuHu reference. The five most highly expressed reliable IFN genes (STAT1, IFITM3, GBP5, GBP2, CXCL10) and the four IFN genes with cross-reactivity ≥ 10% (IRF3, IRF1, IFNB1, DDX58) are labeled. **(c)** Cross-reactivity ratios for the 38 IFN pathway genes, sorted in ascending order. Bars are colored by reliability class: green (reliable, < 10%), orange (moderate, 10–50%), red (unreliable, > 50%). Thirty-four of 38 IFN genes (89%) were classified as reliable. Only three IFN genes exceeded the 50% cross-reactivity threshold (IRF1, IFNB1, DDX58); their exclusion did not alter the overall IFN pathway enrichment conclusions.

**Supplementary Figure 5.**
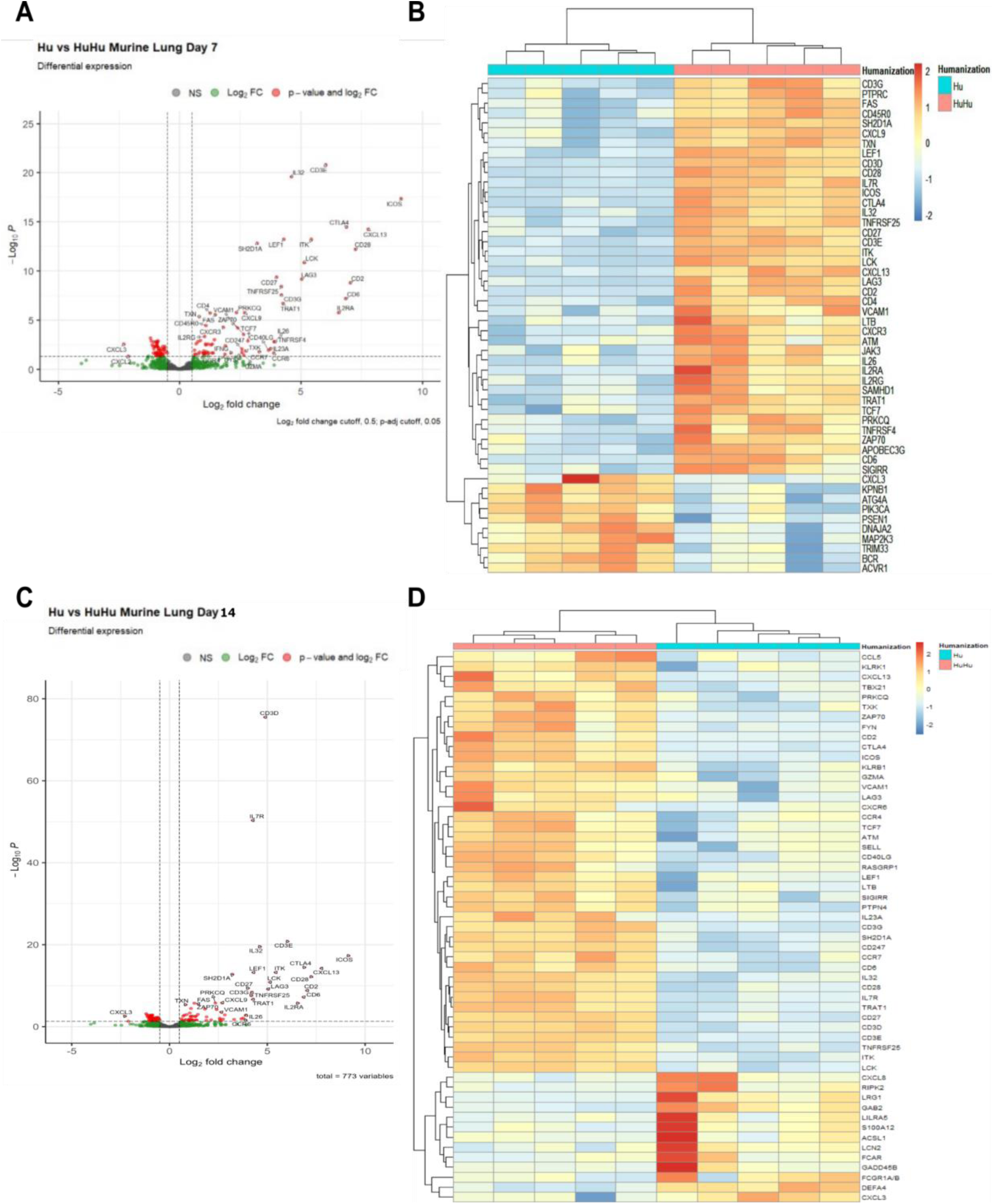
**a.** Volcano plot of differentially expressed genes between infected Hu/Hu and Hu mouse lung samples at day 7 post-infection. **b.** Heatmap of the top 50 differentially expressed genes between Hu/Hu and Hu mouse lung samples at day 7 post-infection. **c.** Volcano plot of differentially expressed genes between infected Hu/Hu and Hu mouse lung samples at day 14 post-infection. **d.** Heatmap of all 55 differentially expressed genes between infected Hu/Hu and Hu mouse lung samples at day 14 post-infection. A *p-adj* cutoff of 0.05 is used for all differentially expressed genes.

**Supplementary Figure 6.**
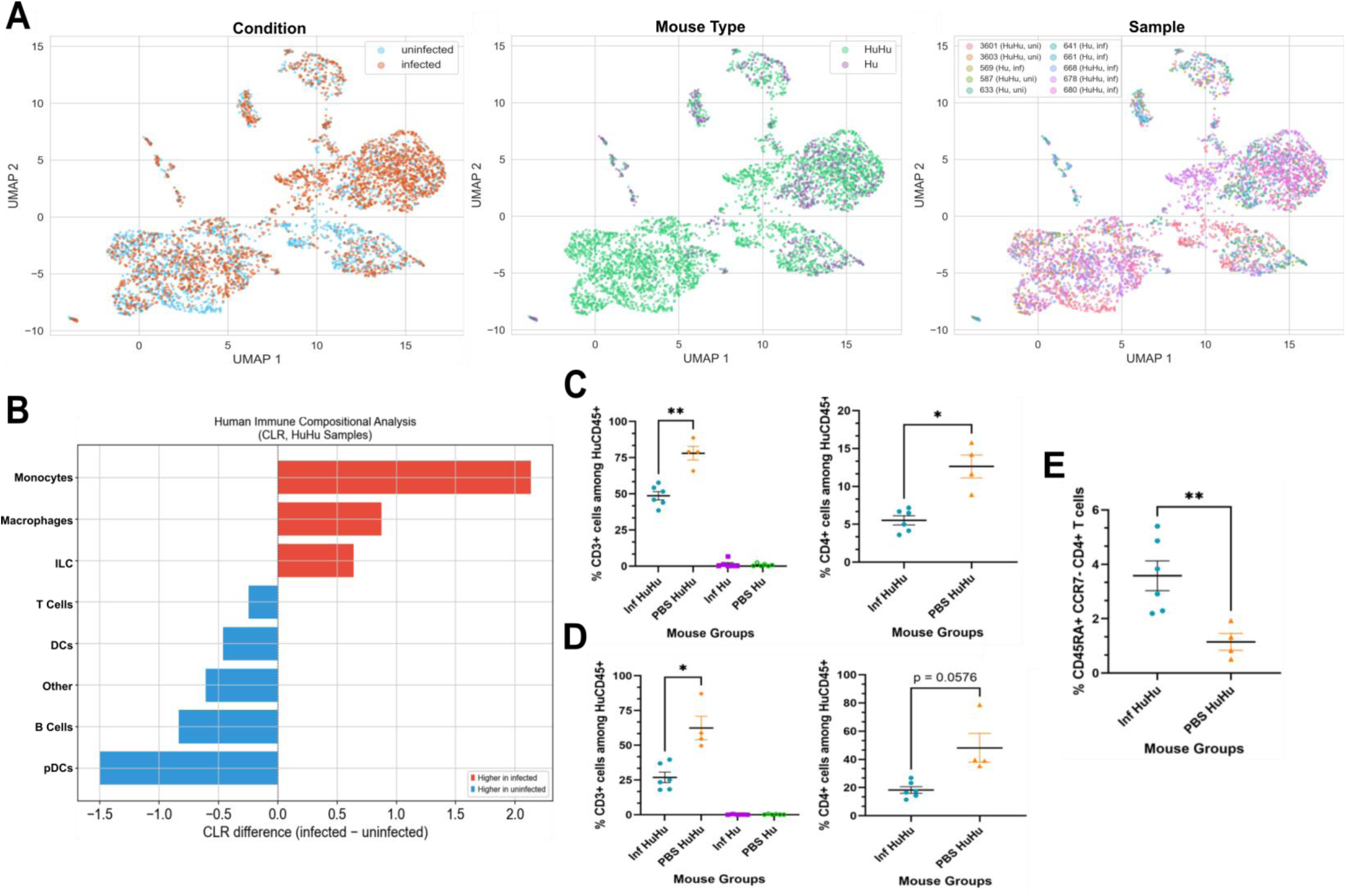
**a.** UMAP of 4,401 human immune lung cells colored by condition (left), Hu/Hu or Hu mouse (center), and sample (right). **b.** Centered log-ratio transformation of human immune cell compositional analysis of Hu/Hu samples in snRNA-seq data. Monocytes CLR p=0.1, Monocytes scCODA inclusion probability = 0.91. **c,d.** Percentage of CD3+ cells (left) and CD4+ cells (right) among all HuCD45+ cells in the mouse lung **(c)** and spleen **(d)** at 8 weeks post-infection detected via flow cytometry analysis. Infected HuHu *n* = 6, PBS HuHu *n* = 4, Infected Hu *n* = 5, PBS Hu *n* = 6. *p-*value of CD3+ cells in the lung = 0.0027 and *p-*value of CD4+ T cells in the lung = 0.0120, *p-*value of CD3+ cells in the spleen = 0.0164. All p-values determined by Welch’s t-test. **e.** Percentage of CD45RA+ CCR7-TEMRA CD4+ T cells in the mouse lung at 8 weeks post-infection detected via flow cytometry analysis. Infected HuHu *n* = 6, PBS HuHu *n* = 4. *p-*value = 0.0056, as determined by Welch’s t test.

**Supplementary Figure 7.**
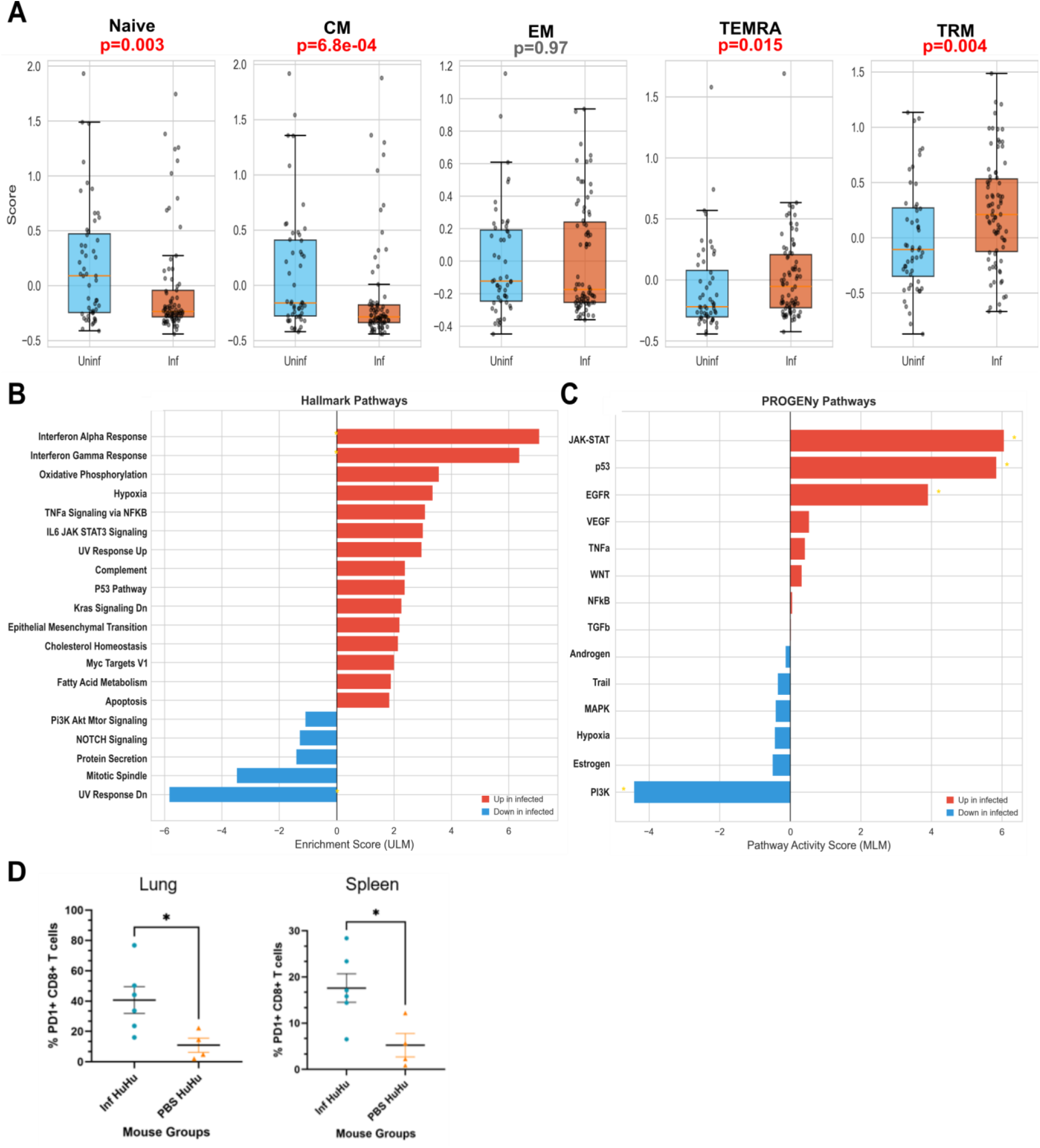
**a.** Gene-set scores computed by ‘scanpy.tl.score\_genes’ for five CD8-relevant subsets: Naive (CCR7, SELL, LEF1, TCF7, CD27, IL7R), central memory/CM (CCR7, SELL, TCF7, IL7R), effector memory/EM (GZMK, CCL5, EOMES, IL7R), TEMRA (CX3CR1, GZMB, PRF1, GZMA, KLRG1, NKG7, GNLY), and tissue-resident memory/TRM (CD69, ITGA1, CXCR6, RUNX3, ITGAE). Box plots show single-cell score distributions; individual cells overlaid as jittered points. *P*-values calculated by Mann-Whitney U test (two-sided). Four of five subsets showed significant differences: Naive (*p* = 3.0 × 10⁻³, down in infected), CM (*p* = 6.8 × 10⁻⁴, down), TEMRA (*p* = 1.5 × 10⁻², up), and TRM (*p* = 4.2 × 10⁻³, up). EM were unchanged. **b.** MSigDB Hallmark pathway enrichment scores computed by univariate linear model (ULM, decoupler) on Wilcoxon rank-sum z-scores for autosomal genes. IFN-alpha response and IFN-gamma response are the top two up-regulated pathways (gold asterisks indicate Bonferroni-adjusted p < 0.05). **c.** PROGENy pathway activity scores computed by multivariate linear model (MLM, decoupler). **d.** Percentage of PD1+ CD8+ T cells in the mouse lung (left) and spleen (right) at 8 weeks post-infection detected via flow cytometry analysis. Infected HuHu *n* = 6, PBS HuHu *n* = 4. *p-*value in lung = 0.0199 and in spleen = 0.0149 as determined by Welch’s t test.

**Supplementary Figure 8.**
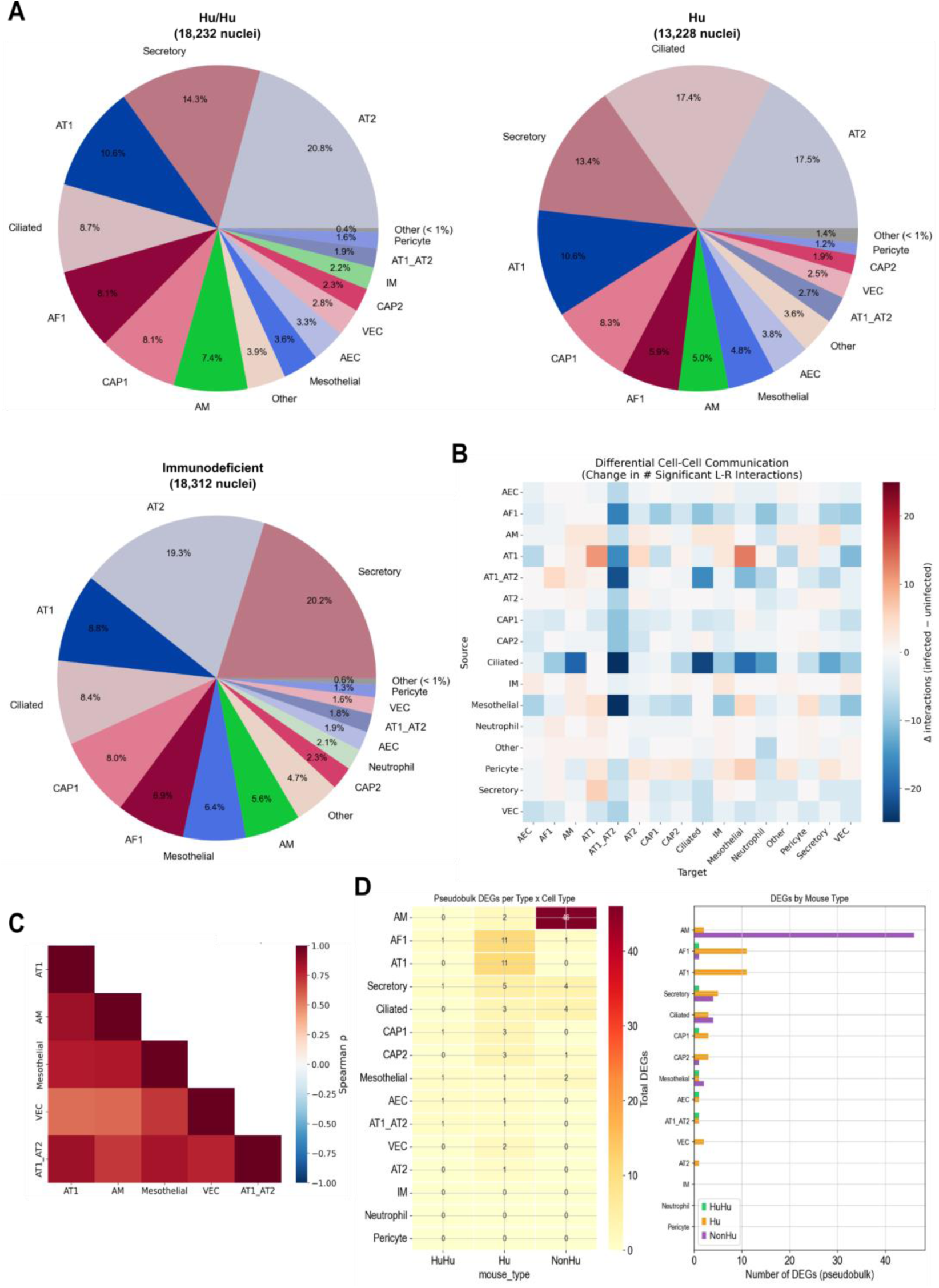
**a.** Pie charts showing the proportion of cell populations in Hu/Hu, Hu and immunodeficient mouse lung samples as determined by clustering of snRNA-seq data. Number of nuclei detected in each group denoted above each pie chart. **b.** LIANA ligand-receptor analysis between infected cell populations. Change in interaction with infected status denoted by heatmap. **c.** Spearman correlation of DE-derived infection response scores across cell types in infected mouse lung. **d.** Pseudobulk DE results stratified by mouse type (HuHu, Hu, NonHu).

**Supplementary Figure 9.**
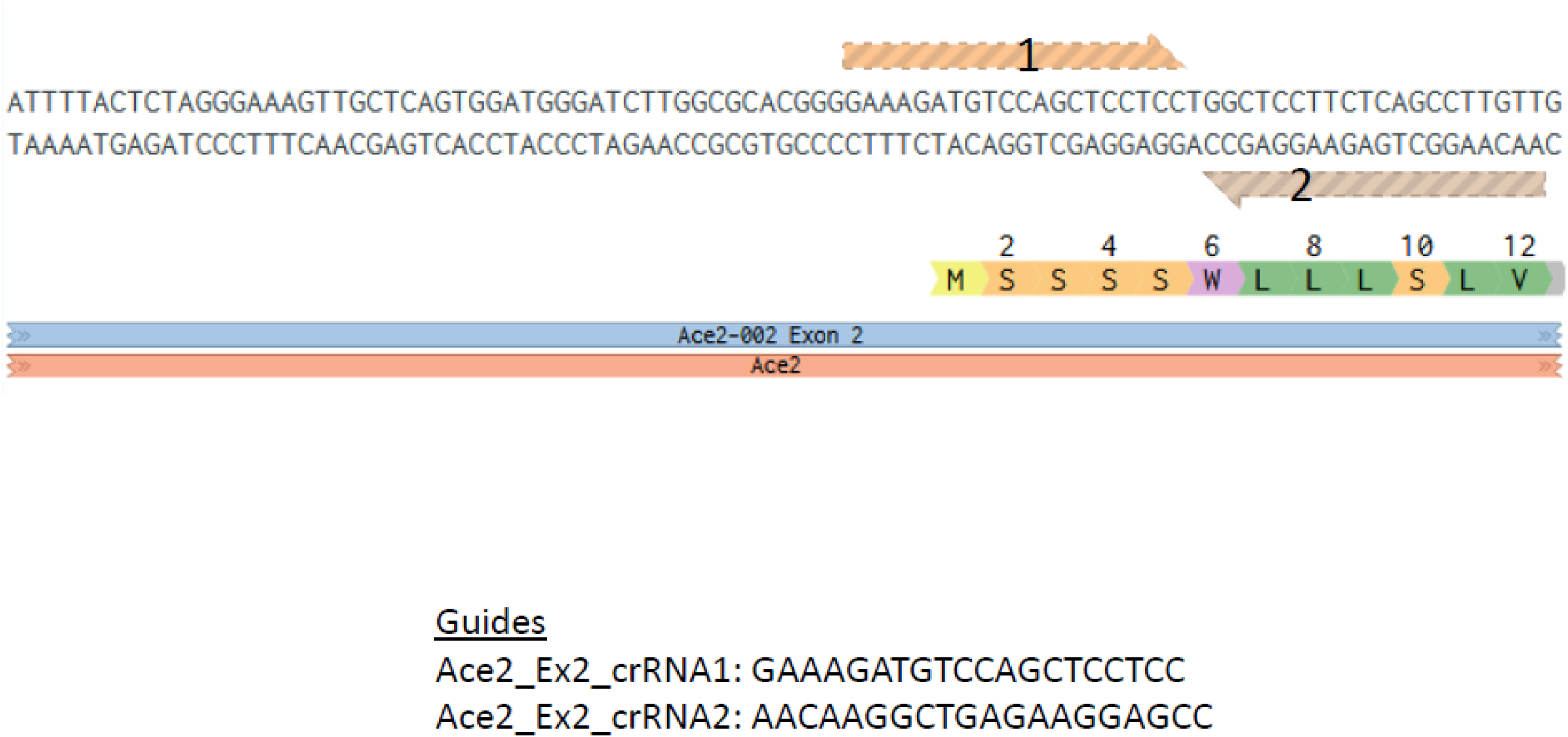
Representative schematic for the insertion of the human ACE2 gene via CRISPR/Cas9 using guide RNAs.

**Table S1:**
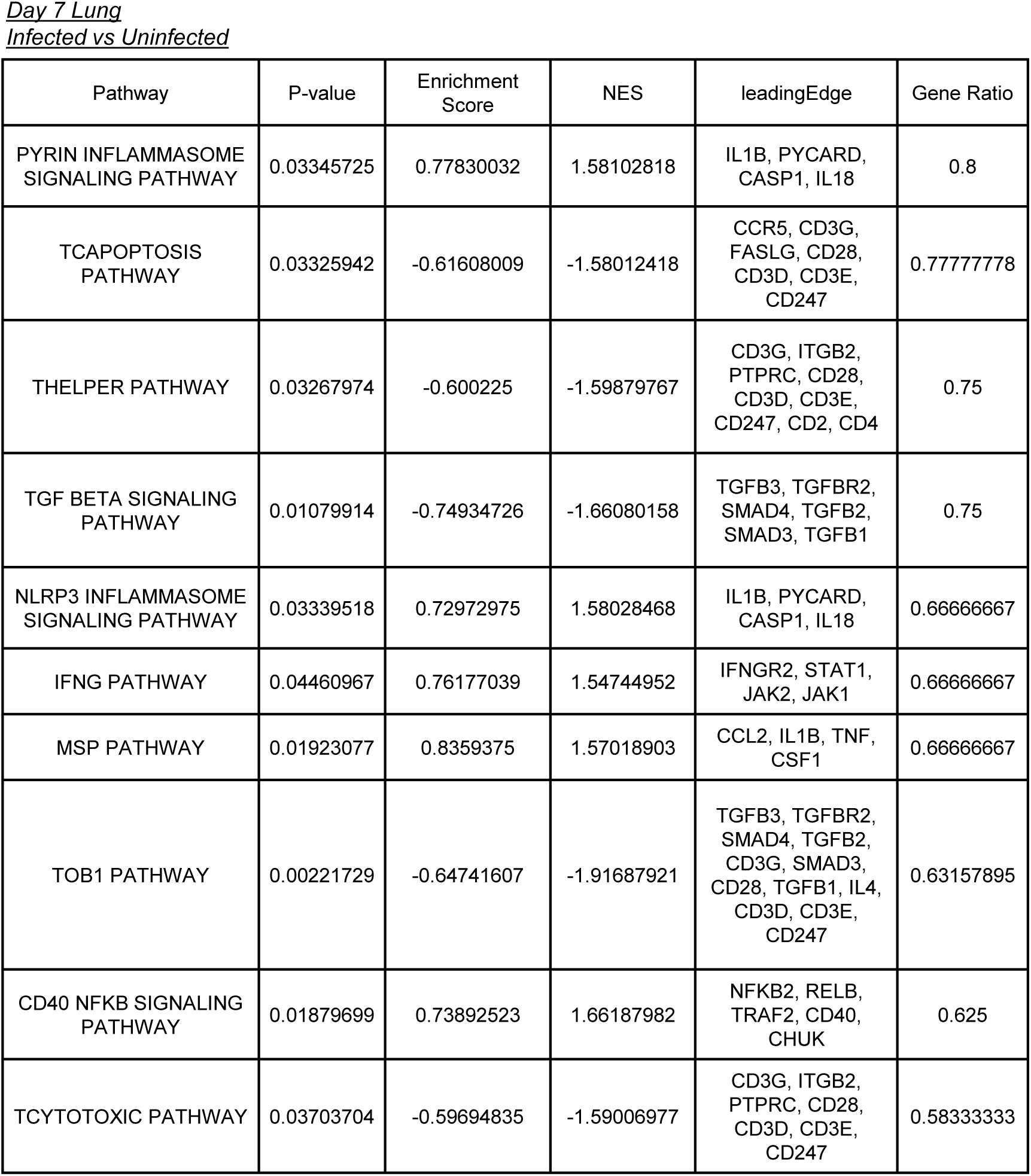
GSEA results for infected and uninfected lung samples at day 7 post-infection.

**Table S2:**
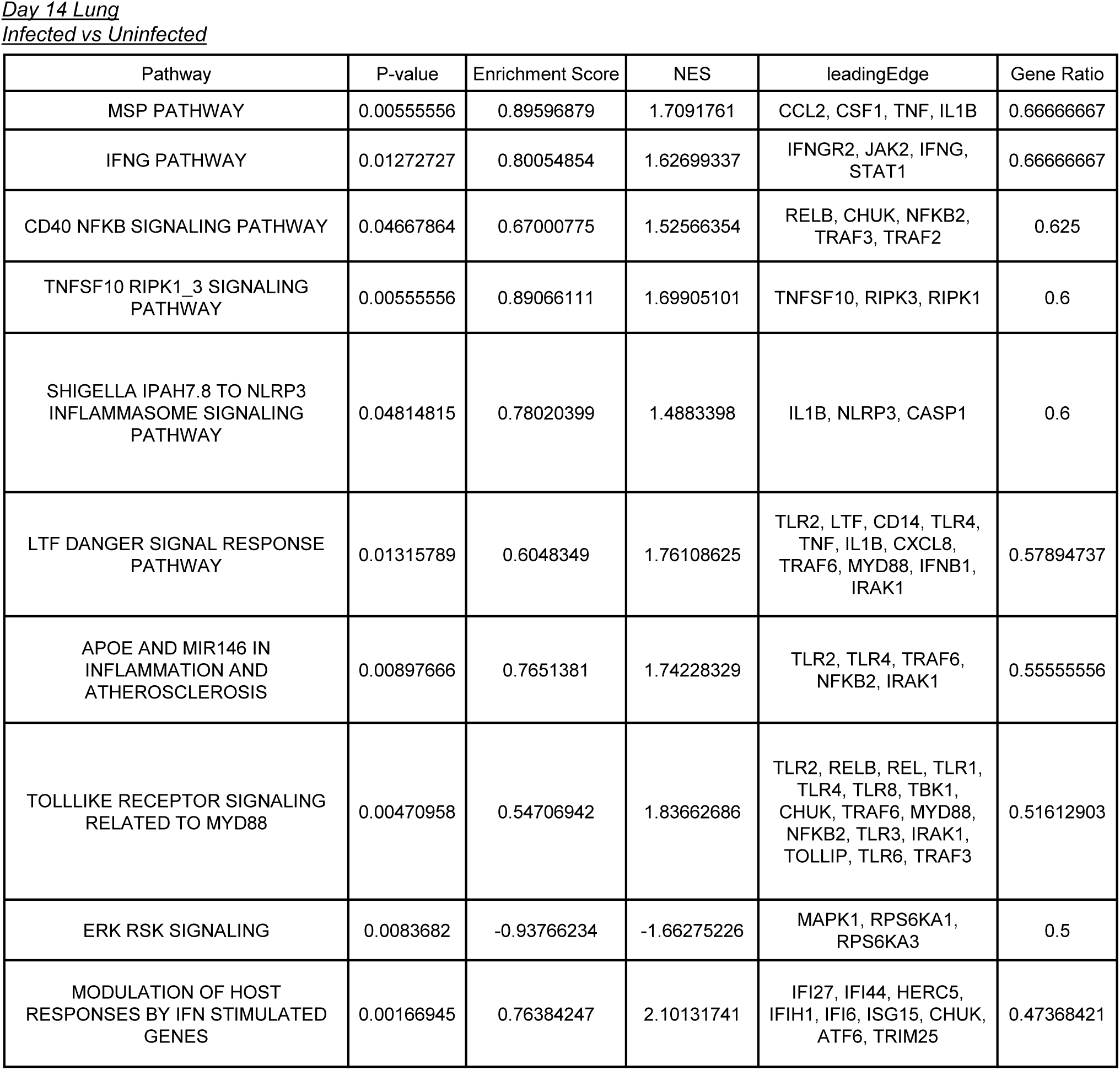
GSEA results for infected and uninfected lung samples at day 14 post-infection.

**Table S3:** GSEA results for infected Hu/Hu and Hu lung samples at day 7 post-infection.

| Pathway | P-value | Enrichment Score | NES | leadingEdge | Gene Ratio |
| --- | --- | --- | --- | --- | --- |
| ALLOGRAFT REJECTION | 0.00119332 | 0.65814998 | 2.27177994 | CD3D, CD3E, CXCL13, ITK, CD28, LCK, CD2, CD3G, TRAT1, IL2RA, CXCL9, CD4, FAS, CXCR3, ZAP70, IL2RG, PTPRC, LTB, CD247, CD40LG, IFNAR2, IFNG, IL16, CD40, CD80, LCP2, IL4R, GZMA, SOCS1, HLA-A, CCR5, IL12B, CCL19, PSMB10, TAP2, CCL5, ITGAL, IL27RA, TAP1, STAT1, IKBKB, WAS, GZMB, HLA-E, IL2, IL2RB, BCL3, GBP2, STAT4, C2, IL10 | 0.255 |
| TGF BETA SIGNALING | 0.00520833 | -0.75674966 | -1.94578969 | ACVR1, TRIM33, THBS1, CDH1, SMAD3, FURIN | 0.11111111 |
| ANDROGEN RESPONSE | 0.01975309 | -0.69939124 | -1.70986152 | HPGD, RPS6KA3, AKT1, MAF, NGLY1, TMPRSS2 | 0.05940594 |
| ANGIOGENESIS | 0.00865801 | -0.92482358 | -1.6043556 | APP, VEGFA | 0.05555556 |
| REACTIVE OXYGEN SPECIES PATHWAY | 0.04513889 | 0.78964129 | 1.52131625 | TXN, SOD1 | 0.04081633 |
| HYPOXIA | 0.00879765 | -0.58574213 | -1.74414219 | VEGFA, PRKCA, TGFB3, DDIT3, BCL2, JUN, RBPJ | 0.035 |
| HEME METABOLISM | 0.02637363 | -0.8342369 | -1.56411563 | MAP2K3, ATG4A | 0.01 |

**Table S4:** GSEA results for infected Hu/Hu and Hu lung samples at day 14 post-infection.

| Pathway | P-value | Enrichment Score | NES | leadingEdge | Gene Ratio |
| --- | --- | --- | --- | --- | --- |
| IL6 JAK STAT3 SIGNALING | 0.04497354 | -0.38068673 | -1.38178716 | EBI3, CXCL3, MAP3K8, CD14, IL1R2, TLR2, CXCL10, IL1R1, CD38, CSF2RB, CCL7, IL6, CSF3R, IL1B, HMOX1, LTBR, IL15RA, CXCL11, MYD88, IRF1, IL17RA, CCR1, SOCS3, TNF, JUN, STAT3, IFNGR2, CD44, TGFB1, IL3RA, CSF1, SOCS1 | 0.36781609 |
| INTERFERON GAMMA RESPONSE | 0.00315457 | -0.41312795 | -1.63334762 | CCL2, SOD2, RIPK2, IFIT2, IFIT3, IFITM3, FPR1, CD274, MT2A, IFI27, CD86, CXCL10, TNFSF10, PTGS2, NAMPT, RSAD2, CD38, CSF2RB, CCL7, CD69, ZBP1, IL6, IFI44, IL15RA, ISG15, CXCL11, MYD88, OASL, IRF1, IL10RA, PELI1, CASP1, TXNIP, IL15, TRIM25, IFIH1, SOCS3, OAS3, GBP4, JAK2, IRF4, STAT3, APOL6 | 0.215 |
| INTERFERON ALPHA RESPONSE | 0.03422983 | -0.41708159 | -1.44439049 | RIPK2, IFIT2, IFIT3, IFITM3, IFI27, CXCL10, RSAD2, IFI44, CCRL2, ISG15, CXCL11, OASL, IRF1, CASP1, TXNIP, IL15, TRIM25, HLA-C, IFIH1, GBP4 | 0.20618557 |
| INFLAMMATORY RESPONSE | 0.01812689 | -0.36682123 | -1.41555341 | CXCL8, CCL2, RIPK2, EBI3, C5AR1, FPR1, CD14, NLRP3, LYN, MARCO, TLR2, OSM, CXCL10, TNFSF10, MEFV, NAMPT, PLAUR, IL1R1, CCL7, CD69, IL6, AHR, IL10, RHOG, ICOSLG, CCRL2, CSF3R, IL18RAP, TLR1, IL1B, IL18, IL15RA, CXCL11, NOD2, IRF1, IL10RA, TLR3, IL15, C3AR1 | 0.195 |
| TNFA SIGNALING VIA NFKB | 0.01139601 | -0.41691043 | -1.57433228 | CCL2, SOD2, RIPK2, IFIT2, GADD45B, CXCL3, CXCL2, MAP3K8, SLC2A3, TLR2, BCL3, CXCL10, PTGS2, NAMPT, PLAUR, PFKFB3, PLEK, BCL6, CD69, IL6, JUNB, ICOSLG, REL, CCRL2, IL1B, IL18, IL15RA, CXCL11, IRF1, IFIH1, SOCS3, MARCKS, TNF, MCL1, JUN | 0.175 |
| ALLOGRAFT REJECTION | 0.00142248 | 0.52202955 | 1.82345775 | CD3D, CD3E, TRAT1, CD3G, ITK, LCK, CD247, CD28, CXCL13, ZAP70, CD2, GZMA, CD40LG, LTB, CCL5, IL2RG, IL2RA, CXCR3, CD4, CD8B, CCR5, IL2RB, PTPRC | 0.115 |
| CHOLESTEROL HOMEOSTASIS | 0.03065134 | -0.80939948 | -1.47448083 | CXCL16, PLAUR, LGALS3, FBXO6 | 0.05405405 |
| WNT BETA CATENIN SIGNALING | 0.04742268 | 0.79202126 | 1.52904894 | LEF1, TCF7 | 0.04761905 |

**Table S5:** GSEA results for infected and uninfected acute heart samples.

| Pathway | P-value | Enrichment Score | NES | leadingEdge | Gene Ratio |
| --- | --- | --- | --- | --- | --- |
| IFNG PATHWAY | 0.03609023 | 0.77719528 | 1.48502176 | JAK2, IFNG, JAK1, STAT1, IFNGR2 | 0.83333333 |
| ICOSLG ICOS PI3K SIGNALING PATHWAY | 0.03618421 | -0.70194733 | -1.72781367 | AKT2, ICOSLG, PIK3CD, AKT1, PIK3R3, ICOS | 0.66666667 |
| TYPE I INTERFERON INDUCTION AND SIGNALING DURING SARSCOV2 INFECTION | 0.0129717 | 0.56596968 | 1.656771 | TBK1, OAS3, TRAF3, EIF2AK2, MAVS, MYD88, OAS2, JAK1, IFNAR1, STAT1, IRF7, IRAK4, TRAF6, TLR7, IFIH1, TLR9, IFNAR2, TLR2, TLR6, TLR4 | 0.64516129 |
| MYD88 DISTINCT INPUTOUTPUT PATHWAY | 0.04198473 | 0.58094519 | 1.5036279 | TIFA, MYD88, TLR5, TRAF6, TLR7, NFKB1, TLR9, TLR2, TLR6, TLR1, TLR4 | 0.61111111 |
| TCAPOPTOSIS PATHWAY | 0.04285714 | -0.61333489 | -1.65885868 | FASLG, CD28, CD4, CD3D, CD3E | 0.55555556 |
| TOLLLIKE RECEPTOR SIGNALING RELATED TO MYD88 | 0.0495283 | 0.49892085 | 1.460498 | REL, TBK1, TRAF3, MYD88, RELB, TLR5, IRF7, IRAK4, CHUK, TRAF6, TLR7, NFKB1, TLR9, TLR2, TLR6, TLR1, TLR4 | 0.5483871 |
| IL10 PATHWAY | 0.01072386 | 0.70190637 | 1.6605533 | IL6, STAT5A, JAK1, IL10RA, HMOX1, TNF, STAT1 | 0.53846154 |
| TLR5 NFKB SIGNALING PATHWAY | 0.03689567 | 0.58978673 | 1.52651197 | IL6, IL12B, MYD88, TNF, TLR5, IRAK4, CHUK, TRAF6, NFKB1 | 0.52941176 |
| REGULATION OF IFNG SIGNALING | 0.03129445 | 0.70446656 | 1.53401529 | JAK2, IFNG, JAK1, PTPN6, STAT1, IFNGR2, SOCS3 | 0.5 |
| ERK RSK SIGNALING | 0.00773196 | -0.95555556 | -1.71909283 | MAPK1, RPS6KA1, RPS6KA3 | 0.5 |

**Table S6:** GSEA results for infected and uninfected long-term heart samples.

| Pathway | P-value | Enrichment Score | NES | leadingEdge | Gene Ratio |
| --- | --- | --- | --- | --- | --- |
| YERSINIA YOPP J TO TLR2 4 MAPK SIGNALING PATHWAY | 0.01724138 | -0.85135135 | -1.79404062 | MAP2K3, MAP2K2, MAP2K4, MAP2K7 | 0.66666667 |
| AML1 ETO FUSION TO PU.1 MEDIATED TRANSCRIPTION | 0.0152207 | 0.90418354 | 1.49472877 | SPI1, ITGAM, CD14 | 0.6 |
| NLRC4 INFLAMMASOME SIGNALING PATHWAY | 0.03929539 | 0.76920069 | 1.45743193 | CASP1, IL1B, NLRC4 | 0.6 |
| MUTATION CAUSED ABERRANT HTT TO TNF JNK SIGNALING PATHWAY | 0.04927536 | -0.85020243 | -1.58425233 | MAPK9, MAPK8, MAP2K7 | 0.5 |
| MUTATION ACTIVATED KRAS NRAS TO ERK SIGNALING PATHWAY | 0.01515152 | -0.78213802 | -1.7872891 | MAP2K2, KRAS, MAPK1, NRAS, RAF1 | 0.5 |
| LTF DANGER SIGNAL RESPONSE PATHWAY | 0.02050114 | 0.62542656 | 1.54534316 | TRAF6, IL1B, TLR2, NFKB1, IRAK4, CD14, TNF, IL6, LTF | 0.47368421 |
| TRK FUSION KINASE TO RAS ERK SIGNALING PATHWAY | 0.01515152 | -0.78213802 | -1.7872891 | MAP2K2, KRAS, MAPK1, NRAS, RAF1 | 0.45454545 |
| UPTAKE AND FUNCTION OF ANTHRAX TOXINS | 0.03333333 | -0.7061793 | -1.72857989 | MAP2K3, MAP2K2, MAP2K4, MAP2K7, CALM1 | 0.45454545 |
| RET FUSION KINASE TO RAS ERK SIGNALING PATHWAY | 0.01515152 | -0.78213802 | -1.7872891 | MAP2K2, KRAS, MAPK1, NRAS, RAF1 | 0.45454545 |
| MACROPHAGE MARKERS | 0.00406504 | 0.86851193 | 1.64560047 | CD163, RAC2, CD14, CD68 | 0.44444444 |

**Table S7:**
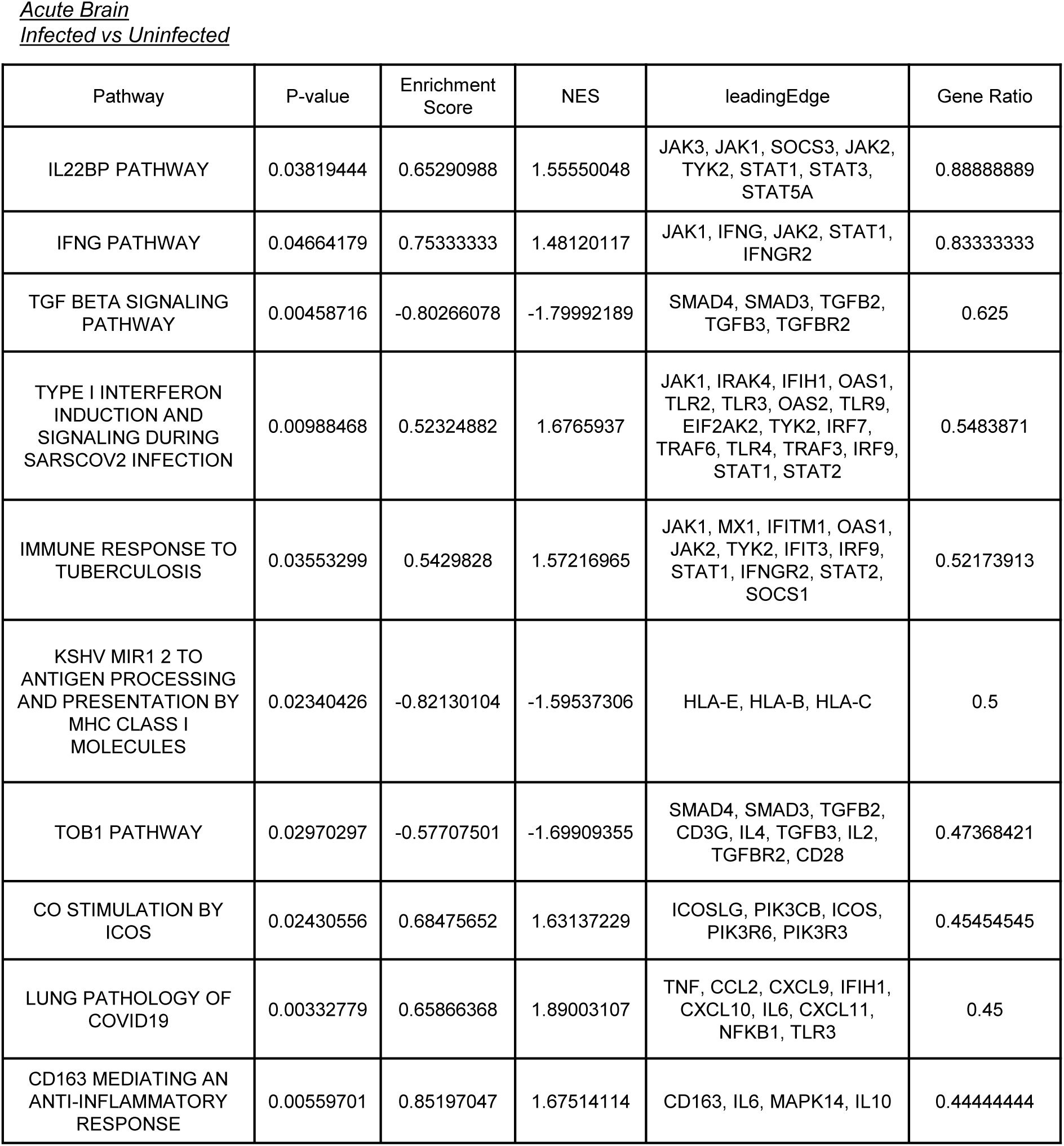
GSEA results for infected and uninfected acute brain samples.

**Table S8:**
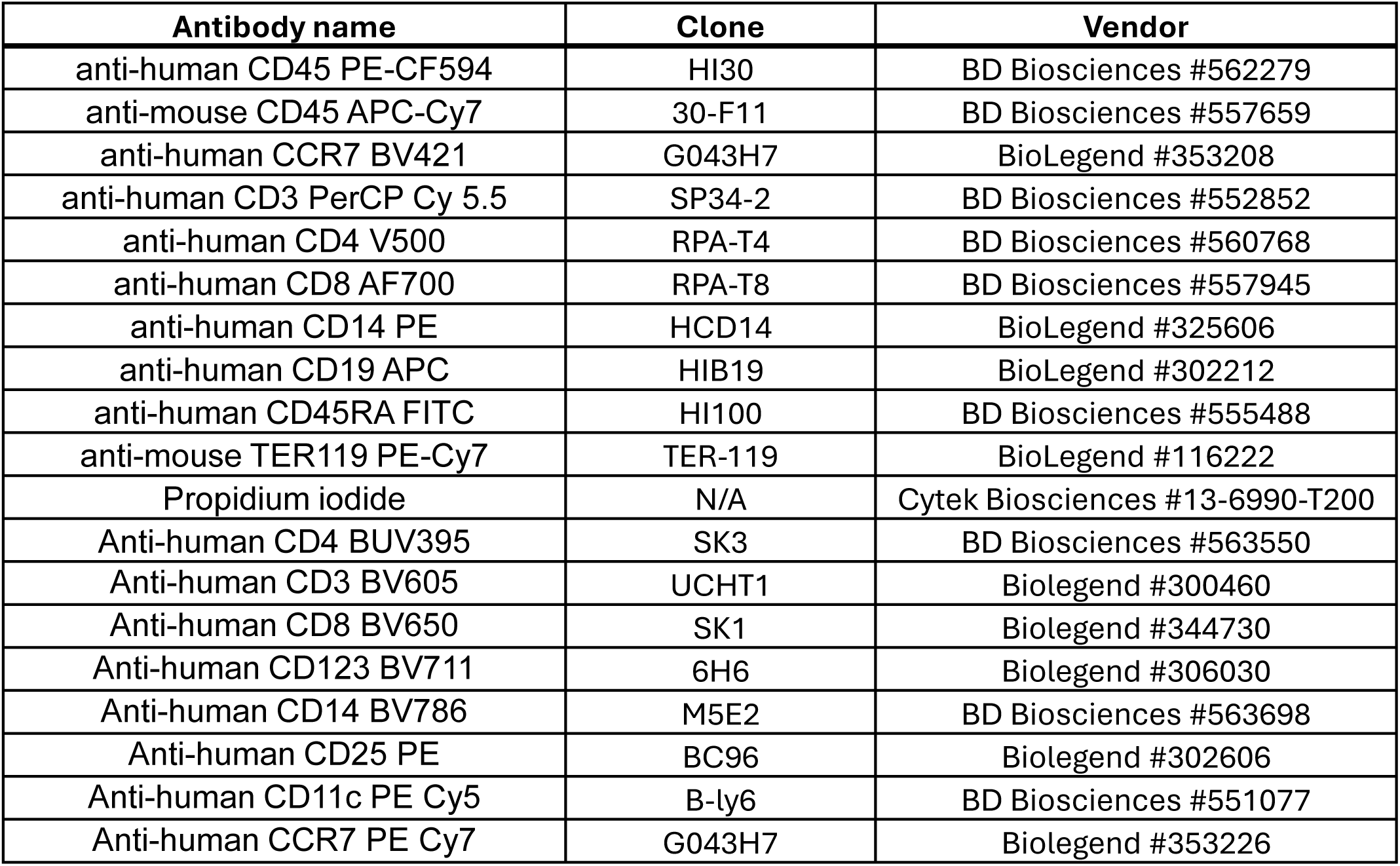
List of antibodies used for flow cytometry analysis.

**Table S9:** List of antibodies used for immunohistochemistry.

| Antibody name | Vendor |
| --- | --- |
| anti-Sars-CoV2-nucleocapsid | Sinobiological #40143 |
| anti-SARS-CoV2 spike glycoprotein | abcam ab272504 |
| anti-hACE2 | abcam ab108209 |

**Table S10:**
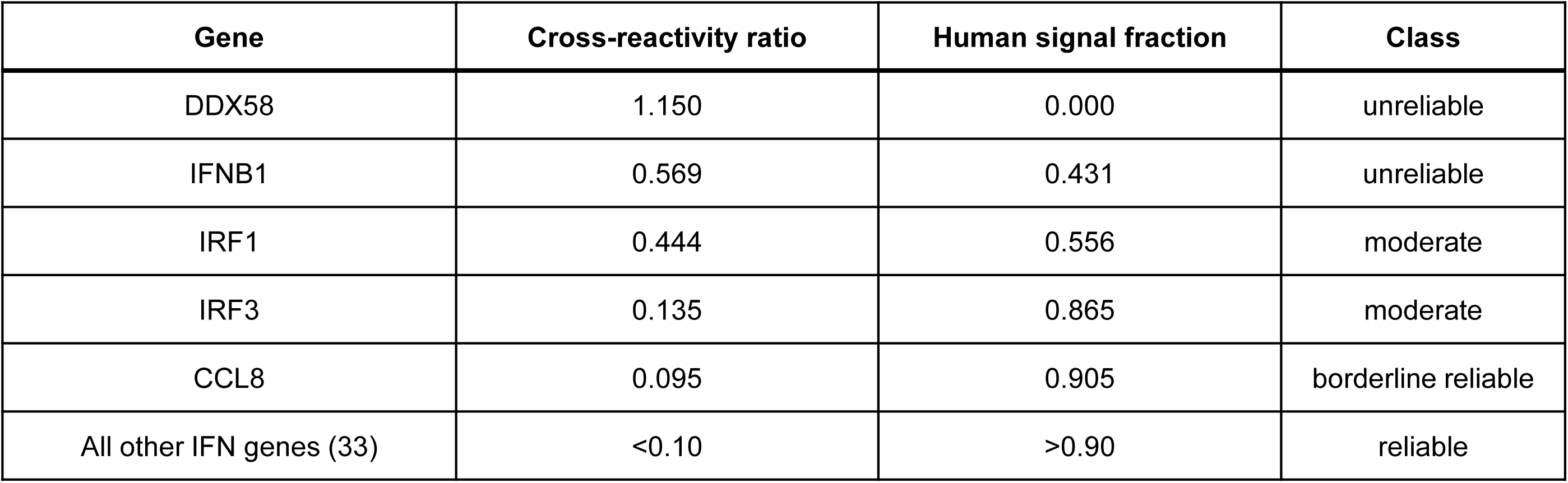
Cross-reactivity of IFN genes.

## Notes

### Competing Interest Statement

The authors have declared no competing interest.

